# Plant pathogenic *Ralstonia* phylotypes evolved divergent respiratory strategies and behaviors to thrive in xylem

**DOI:** 10.1101/2022.11.09.515861

**Authors:** Alicia N. Truchon, Beth L. Dalsing, Devanshi Khokhani, April MacIntyre, Bradon R. McDonald, Florent Ailloud, Jonathan Klassen, Enid T. Gonzalez-Orta, Cameron Currie, Philippe Prior, Tiffany M. Lowe-Power, Caitilyn Allen

**Affiliations:** Department of Plant Pathology, University of Wisconsin-Madison; Microbiology Doctoral Training Program, University of Wisconsin-Madison; Department of Bacteriology, University of Wisconsin-Madison; UMR PVBMT Peuplements Végétaux et Bioagresseurs en Milieu Tropical, CIRAD, France

**Keywords:** Denitrification, denitrifying respiration, vascular wilt, bacterial wilt, endophytic bacteria, niche partitioning

## Abstract

Bacterial pathogens in the *Ralstonia solanacearum* species complex (RSSC) infect the water-transporting xylem vessels of plants, causing bacterial wilt disease. Strains in RSSC phylotypes I and III can reduce nitrate to dinitrogen via complete denitrification. The four-step denitrification pathway enables bacteria to use inorganic nitrogen species as terminal electron acceptors, supporting their growth in oxygen-limited environments like biofilms or plant xylem. Reduction of nitrate, nitrite, and nitric oxide all contribute to virulence of a model phylotype I strain. However, little is known about the physiological role of the last denitrification step, the reduction of nitrous oxide to dinitrogen by NosZ. We found that phylotypes I and III need NosZ for full virulence. However, strains in phylotypes II and IV are highly virulent despite lacking NosZ. The ability to respire by reducing nitrate to nitrous oxide does not greatly enhance growth of phylotype II and IV strains. These partial denitrifying strains reach high cell densities during plant infection and cause typical wilt disease. However, unlike phylotype I and III strains, partial denitrifiers cannot grow well under anaerobic conditions or form thick biofilms in culture or in tomato xylem vessels. Furthermore, aerotaxis assays show that strains from different phylotypes have different oxygen and nitrate preferences. Together, these results indicate that the RSSC contains two subgroups that occupy the same habitat but have evolved divergent energy metabolism strategies to exploit distinct metabolic niches in the xylem.

**IMPORTANCE:** Plant pathogenic *Ralstonia spp*. are a heterogeneous globally distributed group of bacteria that colonize plant xylem vessels. *Ralstonia* cells multiply rapidly in plants and obstruct water transport, causing fatal wilting and serious economic losses of many key food security crops. Virulence of these pathogens depends on their ability to grow to high cell densities in the low-oxygen xylem environment. Plant pathogenic *Ralstonia* can use denitrifying respiration to generate ATP. The last denitrification step, nitrous oxide reduction by NosZ, contributes to energy production and virulence for only one of the three phytopathogenic *Ralstonia* species. These complete denitrifiers form thicker biofilms in culture and in tomato xylem, suggesting they are better adapted to hypoxic niches. Strains with partial denitrification physiology form less biofilm and are more often planktonic. They are nonetheless highly virulent. Thus, these closely related bacteria have adapted their core metabolic functions to exploit distinct micro-niches in the same habitat.

## INTRODUCTION

Bacteria with flexible respiratory metabolisms can grow in environments with fluctuating electron acceptor availability. Oxygen is the most energetically favorable terminal electron acceptor (TEA), but it is not always available to environmental microbes in soil or to pathogenic bacteria in host tissues. Such microbes are often forced to use alternative TEAs (1). Among alternative TEAs, nitrate (NO_3_^-^) is commonly available and yields the most reductive power (2). Nitrate respiration can occur alone when oxygen is limiting, yielding nitrite (NO_2_^-^) and generating a proton motive force to make ATP. It can also be coupled with denitrification, the oxygen-sensitive three-step enzymatic reduction of nitrite to nitric oxide (NO) to nitrous oxide (N_2_O) to dinitrogen gas (N_2_). Separate reductases carry out each step of this pathway. Nitrate-respiring bacteria theoretically gain maximal energy by reducing NO_3_^-^ all the way to N_2_, using all inorganic nitrogen species in this pathway as TEAs.

We previously showed that a plant pathogen, *Ralstonia pseudosolanacearum* GMI1000, depends on nitrate respiration for colonization and virulence (3). GMI1000 is a member of the *Ralstonia solanacearum* species complex (RSSC), a diverse group of bacteria that colonize and block the water-transporting xylem vessels of higher plants (4). Xylem is a relatively low O_2_ environment, containing about 200 μM O_2_, which is hypoxic relative to the 9.4 mM O_2_ in the atmosphere (3). However, xylem sap also contains 30 mM NO_3_^-^, which is the optimal concentration for growth of strain GMI1000 (3). Respiratory reduction of NO_3_^-^, NO_2_^-^;, and NO are all crucial for plant colonization and virulence of GMI1000 (3). However, it has long been known that strains in the RSSC vary in their ability to carry out the final reduction of N_2_O to inert N_2_ gas (5). We wondered whether these closely related bacteria have adapted their core metabolic functions to exploit distinct micro-niches in the same habitat.

The RSSC contains four phylogenetic lineages, phylotypes I-IV (6). Phylotypes I and III are closely related and were recently renamed *R. pseudosolanacearum* while phylotypes II and IV were named *R. solanacearum* and *R. syzygii*, respectively (7). The physiology of the model phyl. I strain GMI1000 has been extensively studied (3, 8–14), but little is known about core metabolism in other plant pathogenic *Ralstonia*. Using a genetically diverse panel of RSSC strains, we showed that phylotypes I and III (hereafter phyl. I/III) are complete denitrifiers that benefit dramatically from the presence of NO_3_^-^during oxygen-limited growth (5). All 25 tested phyl. I/III strains are complete denitrifiers that produce N_2_ gas (15, 16). In contrast, 0/35 phyl. II and 0/8 phyl. IV strains tested produced N_2_ gas (5). Despite this substantial difference in energy metabolism, all RSSC phylotypes include strains that can infect a common host, tomato.

We hypothesized that RSSC phylotypes depend on distinct respiratory mechanisms for growth *in planta*. Using bioinformatics and functional analyses, we identified intriguing metabolic and physiological differences that extend beyond the reduction of N_2_O to N_2_ gas. Phylotype II and IV strains (hereafter phyl. II/IV) lack the final NosZ-dependent denitrification step, and they benefit only slightly from any step in the denitrification pathway under low O_2_ conditions. Intriguingly, phyl. I/III strains and phyl. II/IV strains also differ in biofilm formation, taxis towards low oxygen environments, and aggregation in tomato xylem. These functional differences suggest that despite causing similar wilt disease, these two subgroups have adapted to exploit different respiratory niches *in planta*.

## RESULTS

### Denitrification pathway gene content correlates with RSSC phylotypes

We analyzed the distribution of denitrification genes in 51 genomes reflecting the phylogenomic diversity of the RSSC (Fig. 1A-B). Except for two insect-transmitted phyl. IV strains that have reduced genomes (R24 and BDB R229) (17), all genomes contained the *nar* cluster encoding NarK1/2 nitrate transporters and the first reduction step of the pathway (NO_3_^-^ to NO_2_^-^). All strains also encoded Ani and Nor enzymes that catalyze the second and third denitrification steps (NO_2_^-^ to NO to N_2_O), except for genome-reduced R24. No members of phyl. II/IV carry the *nosZRDFYL* cluster that encodes the terminal N_2_O reduction. In contrast, all phyl. I and 10/11 phyl. III strains have this cluster (Fig. 1C).

**Figure 1.**
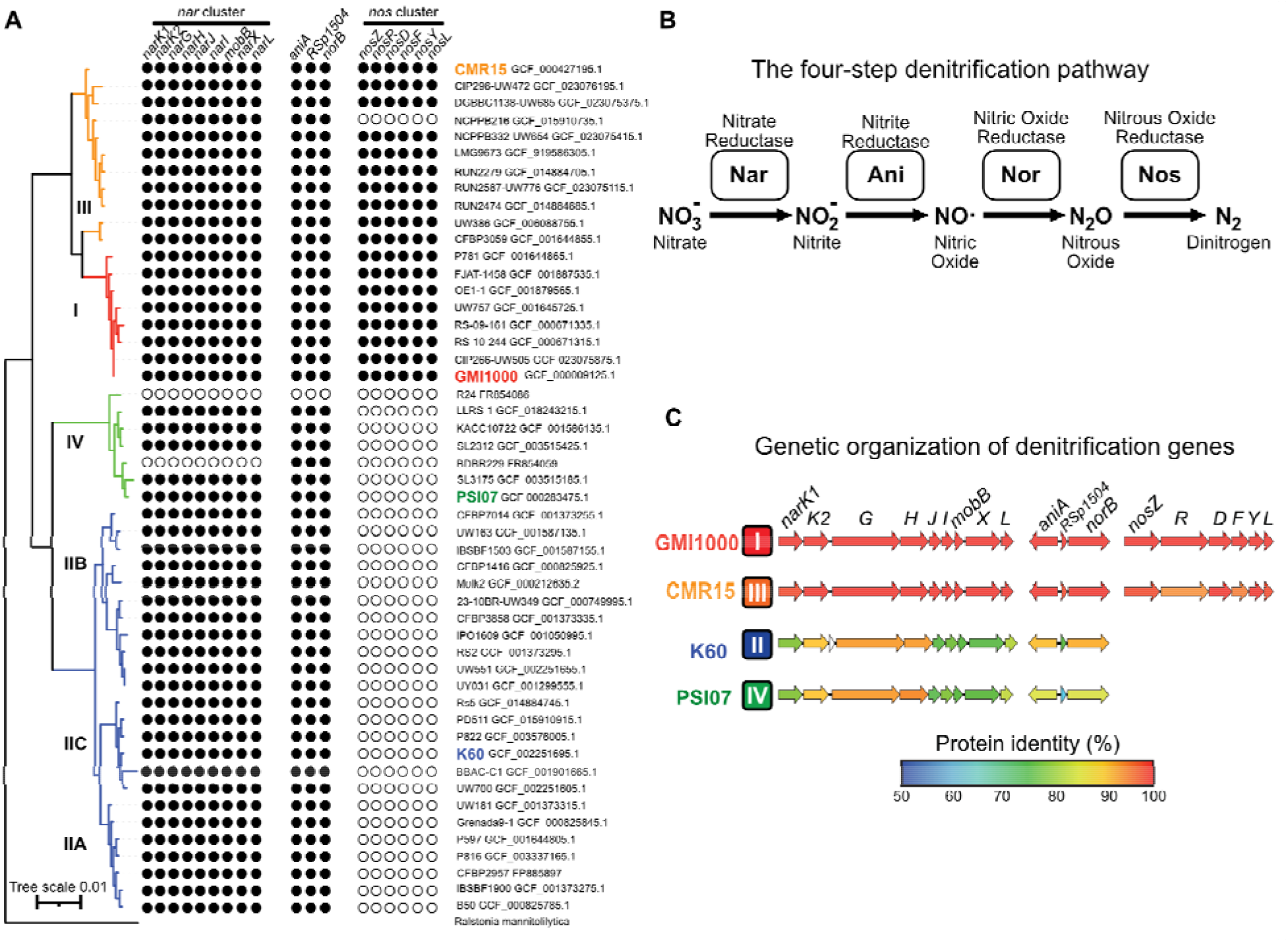
The first three steps of denitrification are well conserved across the RSSC, but only Phyl. I and III have the 6-gene *nos* cluster. **A)** A phylogenetic tree of 51 RSSC genomes that represent the described genomic diversity of the species complex. The tree was constructed in KBase using the Insert genomes into SpeciesTree app. The Newick file was modified in iToL and Affinity Designer. Denitrification genes were identified using the KBase BlastP app with a 40% identity and 70% length cut-off. **B)** The four-step denitrification pathway enzymes, substrates, and products. **C)** Using strain GMI1000 for comparison, the identity values of orthologous proteins were computed with the R package *seqinr* after aligning amino-acid sequences with MUSCLE. Protein percentage identity of each is color coded according to a 50 to 100% gradient. Phylotype is indicated to the right of each strain name. Note that the three gene clusters shown are not adjacent to each other, but they are all located on the megaplasmid.

We wondered if this variation in *nos* content was associated with additional differences in proteins and regulators involved in nitrate-dependent metabolism under low O_2_ conditions. We explored this hypothesis by comparing amino acid sequences of the enzymes, transporters, and regulators encoded in the three denitrification gene clusters. We selected one representative strain from each of the four phylotypes: GMI1000 for phyl. I; K60 for phyl. II; CMR15 for phyl. III, and PSI07 for phyl. IV (Fig. 1). These four strains are all soil-borne, colonize xylem vessels, and cause indistinguishable wilt symptoms on tomato plants. The catalytic subunits NarG, AniA, and NorB were relatively well conserved in all four genomes (Fig. 1C), but the NO_3_^-^transporters (NarK1 and NarK2), the NO_3_^-^-responsive regulators (NarX and NarL), and accessory metabolic proteins (NarJ, NarI, MobB, and RSp1504) had lower sequence identity.

Many highly conserved regulators influence NO_3_^-^metabolism and denitrification in other bacteria (18). All RSSC representatives encoded the potential NO-responsive regulators NnrS (GMI1000 locus RSc3399); NsrR (RSc3397); a predicted O_2_ responsive FNR (RSc1283); two FNR-like regulators (RSp0190 and RSc0966, respectively); and the sigma-54 factor RpoN1 (RSc0408) (19). In contrast, the NorAR NO binding system (RSp0958 and RSp0959) were only present in phyl. I/III (20) (Fig. S1).

### The *nos* N_2_O reduction genes are clustered in an apparently horizontally transferred element scattered among the Beta-proteobacteria

We investigated the presence of the *nosZRDFYL* cluster in complete genomes from diverse strains in the subclass Beta-proteobacteria. As we found in the RSSC (Fig. 1A) and as reported in *Neisseria* spp. (21), the presence of the *nos* cluster was highly variable (Fig. S2).

In genomes of both phyl. I strain GMI1000 and phyl. III strain CMR15, the *nos* clusters are located near fragments of transposition insertion sequences (IS) (Fig. S3). These remnants are phylotype-specific and lie outside the conserved *nos* region. As is typical of horizontally acquired elements, the *nos* cluster is in different locations in the two genomes. This suggests phyl. I/III strains either acquired this locus through separate events or that genome shuffling occurred after the initial incorporation of the cluster.

While the genomic context and the flanking insertion sequences of the *nos* cluster differ between phyl. I/III strains, the cluster gene content and structure is conserved (Fig. S3). The *nos* clusters in GMI1000 and CMR15 contain all eight genes associated with NosZ function *(nosZRDFYLX* and *cco5*) (2, 22–27). Between the IS elements, there are also five conserved genes that are absent from phyl. II/IV genomes, including a SWKP family type III-secreted effector of unknown function (RSp1374).

To gain insight into the evolutionary history of the *nos* cluster, we compared 36 bacterial NosZ protein sequences (Fig. S4). Consistent with multiple horizontal gene transfer (HGT) events, NosZ protein phylogeny generally does not correspond with whole genome phylogeny. Instead, the RSSC NosZ sequences cluster with NosZ from close relatives, *Ralstonia pickettii* and *Cupriavidus metallidurans*. This pattern may indicate a period of vertical inheritance of the cluster followed by its loss in phyl. II/IV, or that HGT is more frequent between organisms with higher sequence homology, independent of transposition events.

### RSSC strains that encode the *nos* cluster require the last step of denitrification for full virulence on tomato

Because all described RSSC strains are plant pathogens, but only two subgroups contain the *nos* locus, we wondered if NosZ contributes to fitness or virulence of phyl. I/III strains. We previously determined that a Δ*nosZ* mutant of phyl. I strain GMI1000 reached a slightly lower population size than wild type in tomato stems 3 days after petiole inoculation (3). However, when we quantified population sizes earlier, at 2 days post inoculation, we found a larger growth difference (Fig. 2A). This early colonization defect suggested that N_2_O reduction contributes to *in planta* success of phyl. I strain GMI1000.

**Figure 2.**
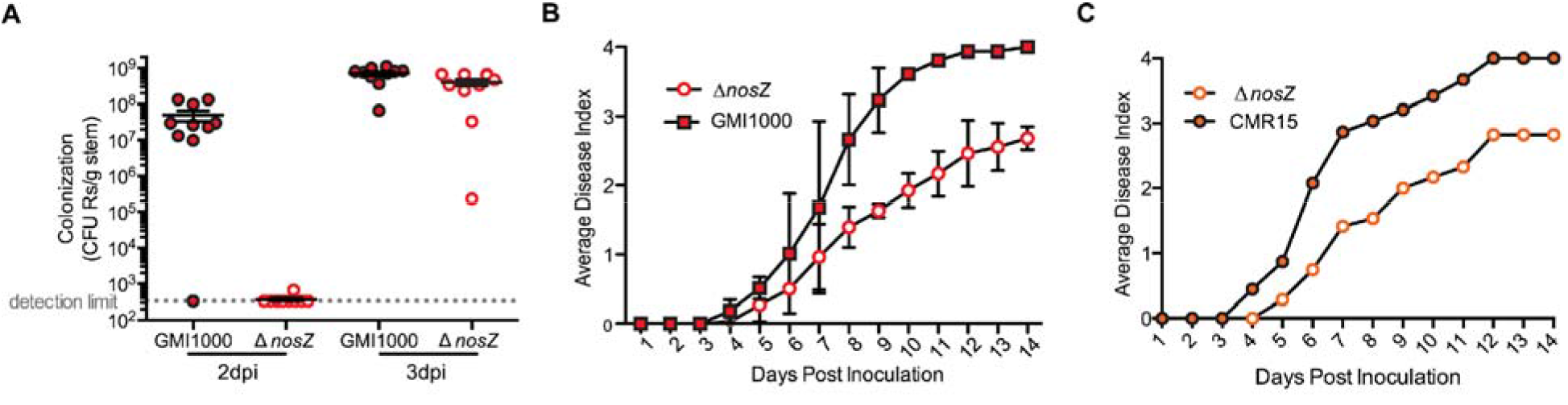
Strains that have *nos* genes require the complete denitrification pathway for full virulence on tomato. **A)** Two and three days after petiole inoculation of 21-day-old wilt-susceptible tomato plants with either wild-type GMI1000 or a Δ*nosZ* deletion mutant, stem sections were harvested and bacterial population sizes in stems were determined by serial dilution plating of ground stem sections. Each circle shows the bacterial population size in a single plant; horizontal bars represent median values; vertical bars indicate standard error of the mean. Data represent ten biological replicates per time point per strain. At both time points, population sizes of GMI1000 and Δ*nosZ* were significantly different (*P*=0.0066 at 2 dpi) and *P*=0.0150 at 3 dpi, two-tailed t-test). The 3 dpi data were published previously ^3^. **B, C)** Symptom severity of RSSC-susceptible tomato plants was monitored daily following naturalistic soil-soak inoculation with 1×10^8^ CFU/g soil of either **B)** wild-type RSSC strain GMI1000 (Phyl. I) and GMI1000 Δ*nosZ* or **C)** wild-type RSSC strain CMR15 (Phyl. III) and CMR15 Δ*nosZ*. Each point indicates average symptom severity; bars in B reflect standard error of the mean of 3 assays, each with 16 plants per treatment (*P*<0.005, 2-way ANOVA). Representative data from one biological replicate containing 12 plants is depicted in panel C.

To investigate the role of N_2_O reductase in virulence for this phyl. I strain, we monitored symptom development following soil-soak inoculations of unwounded plants. This assay revealed that NosZ, and by extension, complete denitrifying respiration, are needed for full virulence in GMI1000 (Fig. 2B). To determine if NosZ is also a virulence factor in the closely related phyl. III, we created a Δ*nosZ* mutant of strain CMR15 and repeated the assay. CMR15 □*nosZ* was similarly reduced in virulence (Fig. 2C). This shows that while strains from all four RSSC phylotypes wilt and kill tomato plants, only phyl. I/III strains require *nosZ* for full virulence.

### NO_3_^-^ enhances anaerobic growth of complete denitrifiers in the RSSC

To test whether RSSC phylotypes differ in their denitrification physiology, we selected three representative strains from each phylotype. Growth assays revealed that the *nos* cluster was required for complete denitrification to N_2_ under anaerobic conditions (Fig. 3A) (5). All tested phyl. I/III strains grew to a 3 to 4-fold higher cell density when provided with NO_3_^-^than when this TEA was absent. In contrast, anaerobic growth of phyl. II/IV strains was unaffected by the presence of NO_3_^-^. Notably, □*nosZ* mutants of both GMI1000 and CMR15 still grew better with NO_3_^-^than without it [data not shown; (3)]. This indicates that the lack of N_2_O reductase activity alone does not explain the inability of phyl. II/IV strains to use NO_3_^-^as a TEA under anaerobic conditions.

**Figure 3.**
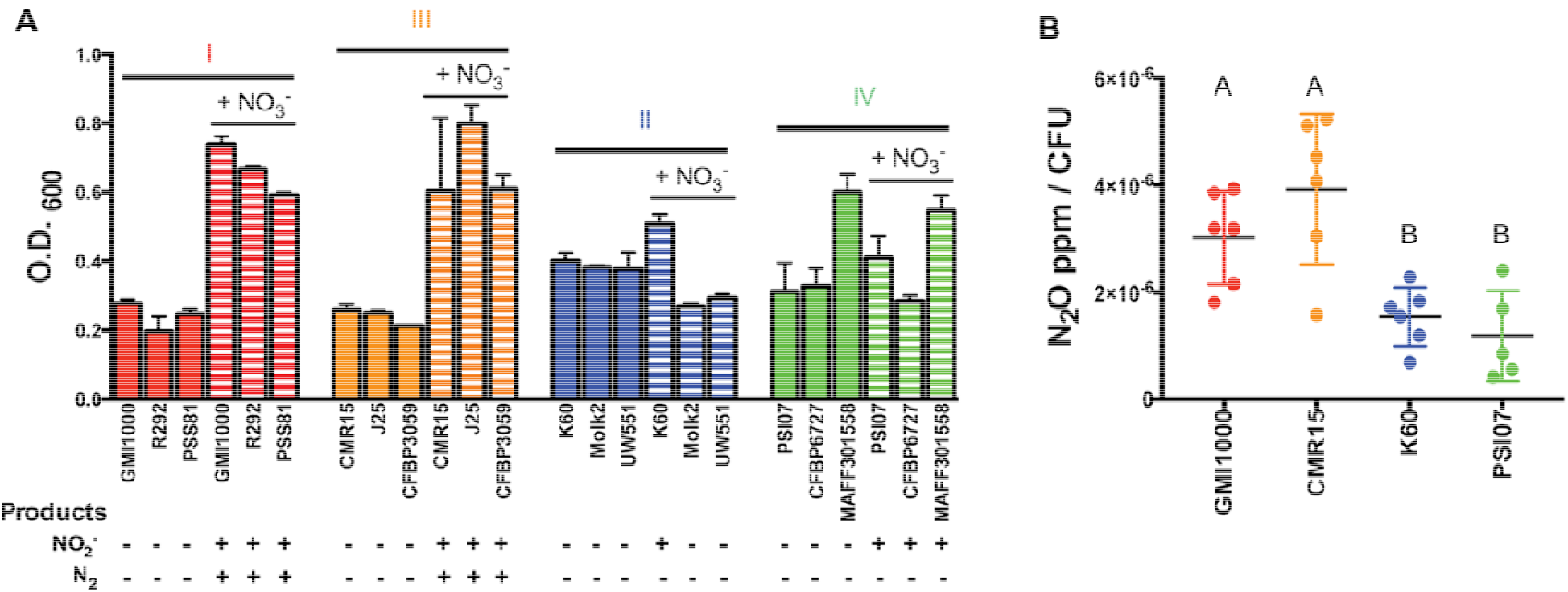
Presence of a functional *nos* cluster in RSSC strains correlates with ability to anaerobically grow on nitrate and produce N_2_O. **A)** Using three representative strains per phylotype (as labeled on X-axis), cell cultures (O.D._600_ 0.001) were incubated statically for 72 hours in VDM with or without 30 mM nitrate and endpoint growth was measured spectrophotometrically. Vertical bars represent standard error of the mean. Growth data were used with permission from ^5^. In separate tubes, Dinitrogen (N_2_) gas production was qualitatively monitored for 96 h in separate tubes. Nitrite (NO_2_^-^) was measured using Greiss reactions, using cultures of each strain in VDM inoculated to O.D._600_ =1.0 and incubated for 3 hours anaerobically. Presence or absence of NO_2_^-^ and N_2_ are indicated with + and -. Data reflect 3 biological replicates per strain. **B)** Production of N_2_O gas. Using a representative strain for each phylotype (GMI1000, K60, CMR15, PSI07), cell cultures (O.D._600_ = 0.0001) were incubated anaerobically for 24 h in VDM with 10 mM NO_3_^-^. CFU was enumerated at time of gas collection. Data shown reflect 5 or 6 biological replicates. Letters indicate *p* <0.05 (Brown-Forsyth and Welch ANOVA with multiple comparisons).

To determine if NO_3_^-^ enhances growth of phyl. II/IV strains at any O_2_ level, we measured growth of each representative strain in 0.1, 1.0, 10.0, and 21.0% O_2_ with or without 30 mM NO_3_^-^ (Fig. 4). At the ambient 21.0% O_2_ all four RSSC strains were inhibited by NO_3_^-^, possibly because of NO_2_^-^ or NO-induced oxidase inhibition (3). Growth of phyl. II K60 was also inhibited by NO_3_^-^ in 10% O_2_. Both phyl. II K60 and phyl. IV PSI07 benefited slightly from the presence of NO_3_^-^ under hypoxic conditions (1.0% or 0.1% O_2_) while phyl. I GMI1000 and phyl. III CMR15 grew much better in this condition. Overall, the phyl. II/IV strains did not grow as well on NO_3_^-^ as phyl. I/III strains at any O_2_ level.

**Figure 4.**
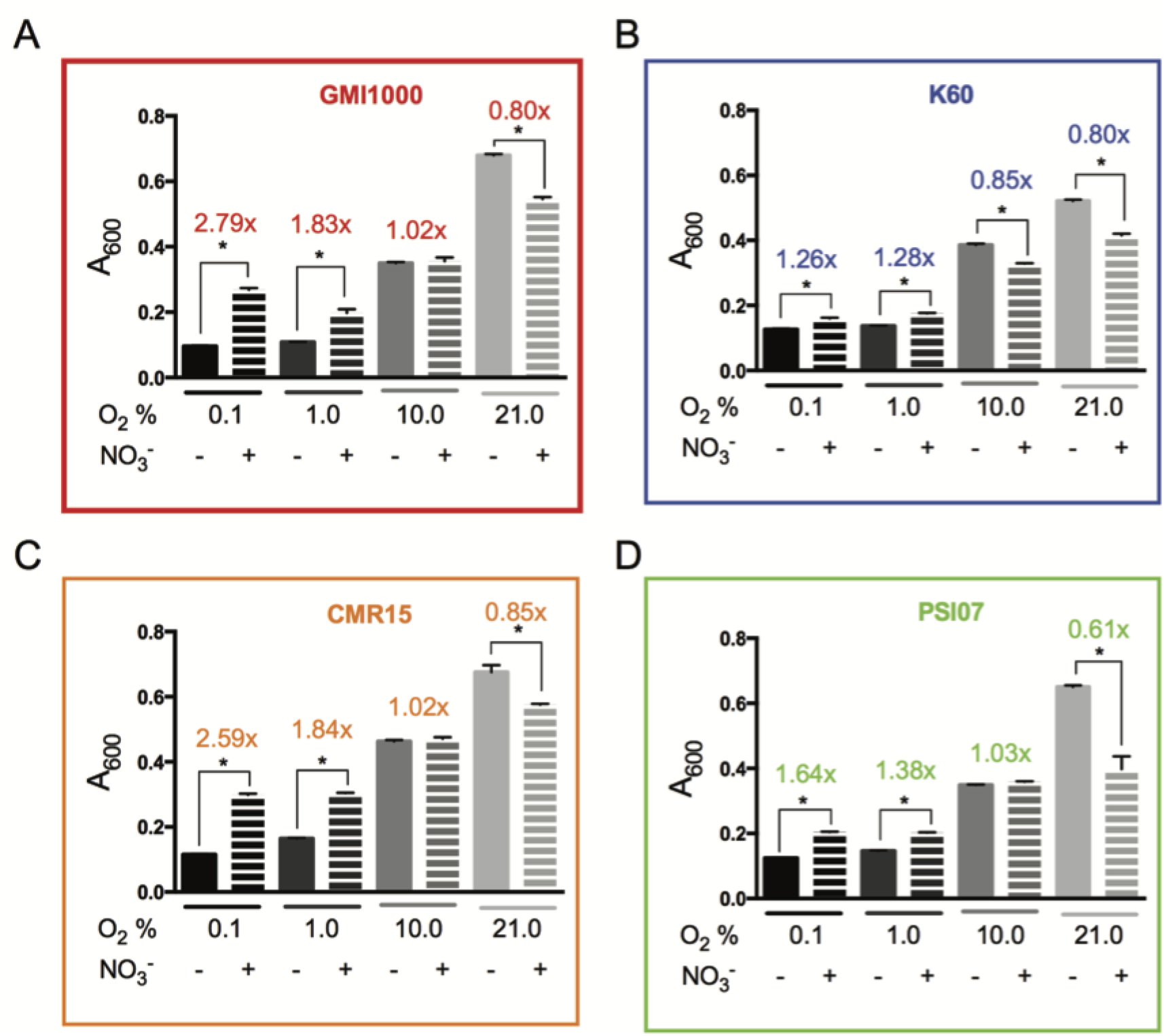
Nitrate enhances growth of RSSC strains under low-oxygen conditions, with the biggest impact on phylotypes I and III. Representative strains of each phylotype (A-D) were grown in denitrifying-favoring VDM broth with and without 30 mM nitrate and at various oxygen levels (0.1, 1.0, 10.0, and 21.0% oxygen). A_600_ was measured after 72 h growth at 28°C, with moderate shaking. Bars indicate standard error. Each treatment was repeated a total of 9 times. \**P*<0.05 by 2-tailed t-test.

All phyl. II/IV strains tested encode the NO_3_^-^, NO_2_^-^, and NO reductases that catalyze the first three steps in denitrifying respiration, but comparative transcriptomic analysis revealed that phyl. II strain UW551 did not express this pathway as highly as phyl. I strain GMI1000 during growth in tomato stems (28). This suggested that denitrification may be less important for phyl. II than for phyl. I. To confirm that the Nar NO_3_^-^ reductases in phyl. II/IV strain are functional, we measured NO_3_^-^ production during anaerobic growth (Fig. 3A). Four of six tested phyl. II/IV strains produced detectable NO_3_^-^ even though their growth was not enhanced by this metabolic conversion. After 4 h incubation at high cell densities (10^9^ CFU/mL), phyl. II strain K60 accumulated ~40 μM NO_2_^-^ while phyl. I GMI1000 accumulated ~90 μM.

To learn if the predicted NO reductase NorB is functional in phyl. II/IV, and to compare N_2_O production by completely and partially denitrifying strains, we measured N_2_O gas produced by denitrifying cultures of the four phylotype representatives. Consistent with the finding that all four strains reduce NO_3_^-^ in low oxygen conditions, all strains made detectable quantities of N_2_O (Fig. 3B). Complete denitrifiers produced approximately twice as much N_2_O per cell than partial denitrifiers (*P*=0.0065).

To determine if phyl. II/IV strains can grow in low O_2_ conditions by using fermentation as an alternative to NO_3_^-^ respiration, we used HPLC to look for fermentation end products in filtered spent culture of the four representative strains after 24 h growth in either aerobic or anaerobic conditions. We did not detect acetate, lactate, or other fermentation end products (data not shown). Moreover, fermentation usually acidifies culture media, and the pH of the RSSC culture supernatants was unaltered.

### Gene enrichment analysis reveals additional metabolic differences between phyl. I/III and phyl. II/IV strains

To identify metabolic functions that may have co-segregated with the *nos* cluster and that could explain the observed differences among strains in inorganic nitrogen respiration, we screened genomes for KEGG categories that are enriched in phyl. I/III strains vs. phyl. II/IV strains (Fig. S5). This analysis suggested that the genomes of complete and partial denitrifying strains are enriched in distinct sets of aromatic degradation pathways. Phyl. II/IV strains were enriched in a Ben/Cat pathway for benzoate/catechol degradation (29). Phyl. I/III strains showed enrichment of a partial Dmp pathway that is missing the DmpB gene that catalyzes the ring opening of catechol (30). The KEGG enrichment analysis was performed on the small set of genomes available in 2015, which was biased towards phyl. II genomes. We curated high-quality genomes that better represent the genomic diversity of the RSSC and performed BlastP searches for aromatic degradation enzymes (Fig. 5). This robust analysis confirms that the Ben/Cat pathway is indeed present in most phyl. II/IV genomes. The partial Dmp pathway is sporadically present in phyl. I/III genomes and absent from phyl. II/IV genomes. Overall, the protocatechuate degradation pathway (Pca) (29) is broadly conserved across the RSSC, as we previously found (31). The hydroxycinnamic acid degradation pathway (Fcs) (31) is broadly conserved except in the phyl. IIC lineage and the Banana Blood Disease lineage of genome-reduced phyl. IV strains. The salicylic acid degradation pathway (Nag) (32) is broadly conserved except in phyl. IIC and most IIA genomes and the Sumatra Disease of Clove lineage of genome-reduced phyl. IV strains.

**Figure 5.**
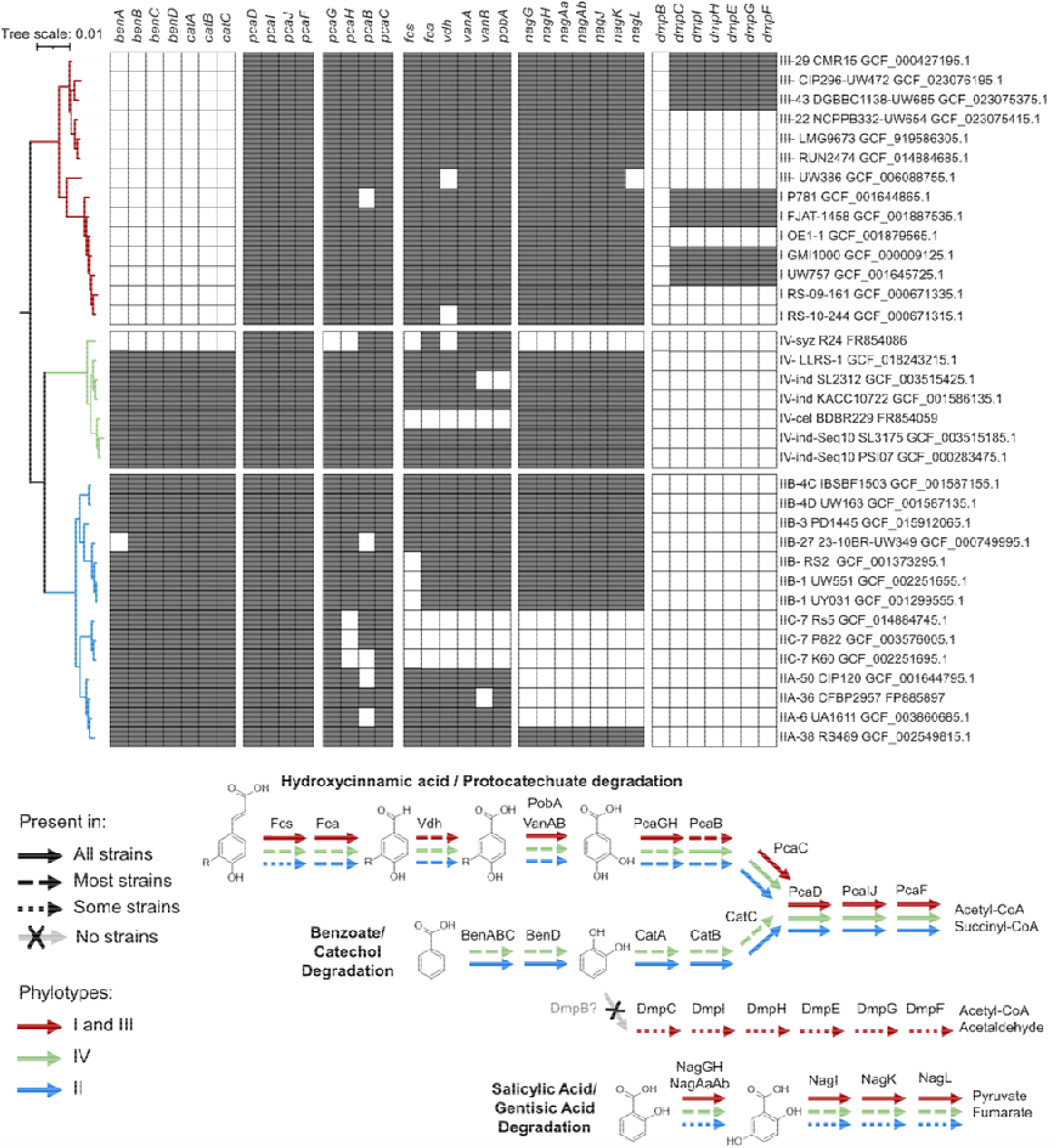
Variation in aromatic degradation pathways within the RSSC. Top: presence/absence of aromatic degradation genes across the species complex. Bottom: Summary of the conservation of the aromatic degradation pathways across distinct phylotypes. The benzoate/catechol degradation pathway and the hydroxycinnamic acid/protocatechuate pathways converge on the last four Pca enzymes. The phylogenetic tree was constructed based on 49 conserved genes using the KBase SpeciesTree app. The presence/absence of each gene was determined through BlastP analysis of the genomes.

### Complete and partial denitrifiers in the RSSC have different oxygen preferences

To better understand oxygen preferences within the RSSC, we stab-inoculated strains into tubes of VDM soft agar with or without 30 mM NO_3_^-^. In these tubes, O_2_ is available in a diffusion gradient near the agar surface. After one week, the growth patterns in these culture tubes indicated that phyl. I/III GMI1000 and CMR15 had a strong preference for anaerobic environments when NO_3_^-^was available, while phyl. IV had a subtle preference for lower-than-atmospheric O_2_ levels (Fig. S6). A GMI1000 Δ*narG* mutant, which lacks the first denitrification step, had no tactic response to NO_3_^-^conditions (data not shown), which is consistent with dependence of these migration patterns on energy taxis (also known as aerotaxis) (33).

### Complete and partial denitrifiers in the RSSC may have adapted to different ecological niches

Biofilms are typically hypoxic (34), so we hypothesized that phyl. I and III strains would form more robust biofilms. We tested this hypothesis using the PVC-crystal violet biofilm assay, which showed that phyl. I/III strains formed thicker biofilms *in vitro* than phyl. II/IV (Fig. 6B).

**Figure 6.**
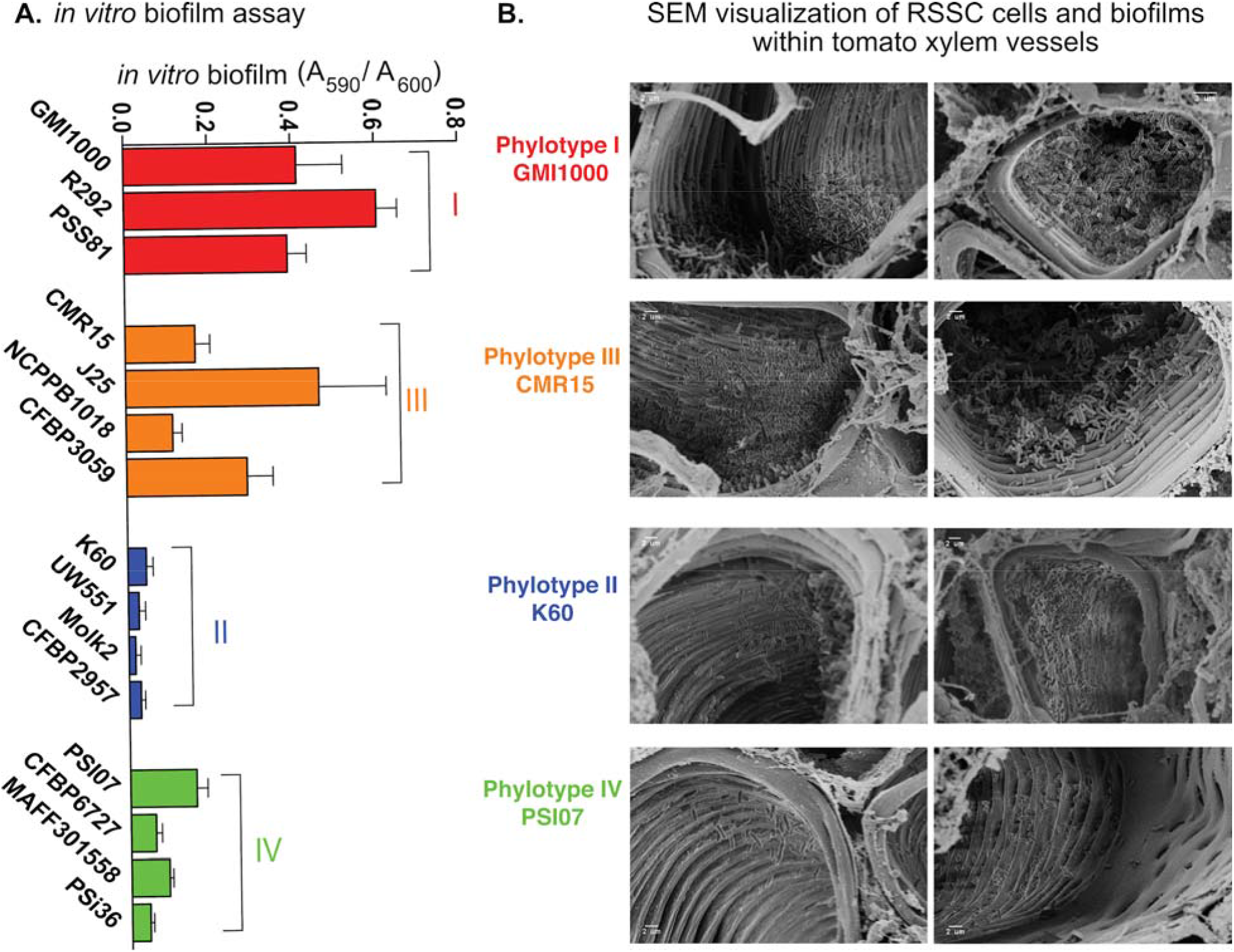
Complete denitrifying RSSC strains form more biofilm in culture and occupy different niches in tomato xylem during bacterial wilt disease. **A)** Using 3-4 representative strains per phylotype, biofilm formation was quantified using the PVC plate-crystal violet assay 96 well plates. Vertical bars represent standard error of 9 biological replicates. **B)** Representative SEM images showing stem cross-sections of tomato plants infected with a representative strain from each phylotype (GMI1000, K60, CMR15, PSI07), as indicated. Susceptible tomato plants (cv. Bonny Best) were soil-soak inoculated with ~1×10^8^ CFU/mL bacterial suspension in 80 g soil. At the first sign of disease, stem samples were taken, sliced, fixed, plated with gold, and visualized with a Zeiss LEO 1530 high-resolution scanning electron microscope. SEM images were captured from two biological replicates of plant inoculations each, with three or four plants and two stem slices per plant.

Previous SEM studies showed that the complete denitrifier GMI1000 forms dense biofilms in tomato xylem vessels (35, 36). To determine if all RSSC phylotypes form similarly dense biofilm *in planta*, we used SEM to image xylem tissue at onset of wilt symptoms. The complete denitrifying strains in phyl. I/III colonized xylem vessels differently than the phyl. II/IV partial denitrifiers. Phyl. I GMI1000 and phyl. III CMR15 colonized many xylem vessels extensively and often formed thick biofilms on vessel walls and in the lumens (Fig. 6A). In contrast, cells of phyl. II K60 and phyl. IV PSI07 were visible in fewer xylem vessels and often formed single-cell layers on the vessel walls (Fig. 6A). During SEM sample preparation, stem cross-sections are washed in a fixative solution. We hypothesized that planktonic or loosely attached cells may disperse into the solution before the fixative can anchor them in place. To assess the relative numbers of planktonic and attached cells in plants infected with each strain, we quantified bacterial populations in homogenized stem samples and then quantified the unattached cells that streamed from cut stem sections incubated in water. As previously observed, all four phylotype representatives colonized plants similarly, reaching population sizes >1×10^9^ CFU/g stem (Fig. S7A). There was no difference in the proportion of released cells of phyl I/III strains GMI1000 and CMR15 or of phyl. II K60. More than 90% of these cells remained in cut stem sections. However, almost twice as many phyl. IV PSI07 cells streamed from cut stems into the water (Fig. S7B). If PSI07 cells were more often planktonic or loosely attached, this could explain the relatively few bacteria visible in SEMs of PSI07-infected stems. Together, the PVC biofilms, *in planta* SEM images, and streaming assay results suggest that during plant infection, complete denitrification may help phyl. I/III strains form more robust biofilms on xylem walls.

To assess the effects of bacterial growth on O_2_ levels in tomato xylem, we used a microprobe to directly measure O_2_ concentrations in sap from plants colonized by each of the four representative strains. Bacterial wilt disease significantly reduced O_2_ availability in xylem sap regardless of the infecting RSSC strain. This finding is consistent with our previous observation that infection by GMI1000 reduced xylem O_2_ levels by half relative to healthy plants (3). The O_2_ levels in stagnant water and xylem sap from healthy tomato plants are 230 μM O_2_/L, and around 200 μM O_2_/L, respectively (Fig. 7). Sap from early to mid-stage diseased plants (disease index = 1-3) infected with any of the four representative RSSC strains contained 220 to 95 μM O_2_/L. Oxygen levels were lower in sap from late-stage diseased plants, ranging from 175 to 0 μM O_2_/L. There was little difference in sap oxygen levels between tested strains (Fig. 7). This suggests that phyl. II/IV and I/III strains all rapidly consume any available oxygen and thus experience similarly low O_2_ during wilt pathogenesis. Nonetheless, our genomic and functional analyses indicate that the two groups have adapted to this metabolic challenge in distinct ways.

**Figure 7.**
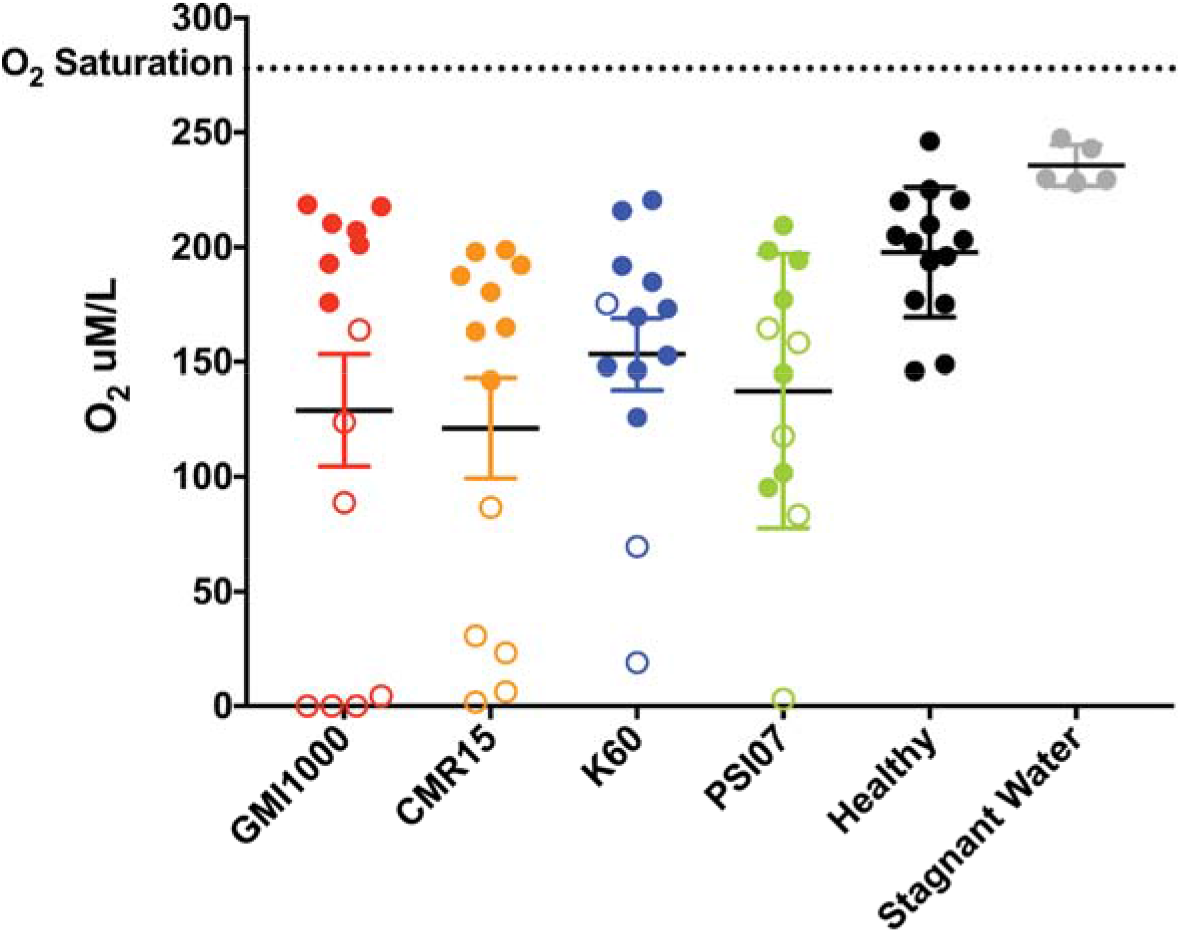
Xylem sap oxygen levels in healthy and RSSC-infected plants. A microsensor was used to measure oxygen levels in the xylem sap that exuded from a freshly-detopped tomato plant inoculated with water (Healthy) or with RSSC phylotype representatives, GMI1000, CMR15, K60, or PSI07. Each circle represents the O_2_ level in one plant; N =12-14 plants/condition. Closed circles are levels from plants with mid-stage disease (1-74% leaf area wilted) and open circles are levels from plants with late stage disease (>75% leaf area wilted). Xylem sap from infected plants contained less O_2_ than sap from healthy plants (*P*= 0.0443, 0.0015, 0.0125, and 0.0031 for healthy vs. GMI1000, CMR15, K60, and PSI07, respectively, Mann-Whitney test). All tested RSSC strains similarly reduced sap O_2_ levels. Xylem sap from healthy plants contained less O_2_ than stagnant water (*P*= 0.0005, Mann-Whitney test). Bars show standard error of the mean.

## DISCUSSION

We discovered that the four phylotypes in the RSSC have surprisingly diverse energy metabolisms. Comparative genomics of diverse RSSC strains revealed that the first three steps of denitrification are broadly conserved across the RSSC, but only phyl. I/III have the final Nos-dependent step. These bioinformatic results are consistent with prior physiological studies showing that only phyl. I/III strains produce N_2_ gas (5). Unfortunately, this means that the official taxonomic revision of RSSC species incorrectly describes the denitrification phenotypes of *R. pseudosolanacearum* and *R. solanacearum* (7). This warrants correction.

Plant xylem has often been described as a nutrient-poor environment (9). However, recent quantitative metabolomics and physiological modeling shows that healthy tomato xylem sap contains enough organic carbon and nitrogen to sustain *Ralstonia’*s rapid growth, including mM levels of amino acids and 0.1 mM levels of glucose and sucrose (12, 13). Xylem sap should thus be considered a rich, but incomplete, medium (37). *In vitro* experiments indicate that phyl. I strain GMI1000 relies primarily on the glutamate, asparagine, aspartate, and glucose in xylem (12, 13). Catabolism of other sugars contribute slightly to success in xylem (11). Although GMI1000 can assimilate NO_3_^-^ into organic nitrogen via NasA, NO_3_^-^ assimilation appears to be most important for *Ralstonia* success in the rhizosphere at the earliest early stage of disease (38). Our data indicate that for phyl. I/III strains, NO_3_^-^ is primarily a terminal electron acceptor *in planta* rather than a nitrogenous building block.

If phyl. II/ IV strains lack the Nos enzymes, then why do phyl. I/III strains require *nosZ* for full virulence on tomato? Pathway blockage could cause the Δ*nosZ* mutants of GMI1000 and CMR15 to accumulate N_2_O. Although N_2_O has low toxicity, these mutants may experience inhibitory N_2_O levels *in planta*. However, direct measurements found that these mutants did not accumulate more N_2_O than wildtype *in vitro*. Because all strains produce similar N_2_O levels during anaerobic growth, the most likely explanation is that phyl. I/III strains have become dependent on Nos enzymes to generate a sufficient proton motive force to support growth *in planta*. Recently, we showed that levels of the denitrification intermediate NO influence expression of the type 3 secretion system for phyl. I strain GMI1000 (39). The type 3 secretion system is a major virulence factor. It is possible that *nosZ* mutants have slightly reduced expression of type 3 secretion system genes.

Throughout plant infection, RSSC pathogens experience dynamic levels of O_2_ and reactive oxygen/nitrogen species. We found that infection by all representative strains similarly decreased O_2_ levels in tomato xylem sap, leading to anoxia in advanced wilt disease. Nevertheless, diverse RSSC strains require mechanisms to manage the reactive oxygen and nitrogen species during plant infections (20, 40–42). Although O_2_ levels decrease during bacterial wilt, some O_2_ is present when *Ralstonia* populations are growing explosivly during early and mid-stage disease. Previously, we showed that a different phyl. II strain (UW551) requires a high affinity cbb_3_-type cytochrome C oxidase for maximal growth *in planta* and in microaerophilic culture (43). Genomes from representatives of all RSSC phylotypes encode both major types of terminal oxidase families: proton pumping heme-copper oxidases and non-pumping *bd*-type oxidases (data not shown; (44)). Nevertheless, it remains unclear if the two groups of RSSC strains vary in their ability to use oxygen as a TEA.

Because phyl. I/III strains form abundant and thick biofilms on xylem vessel walls, these strains may experience much lower O_2_ levels *in planta* than were reflected in our measurements. Although O_2_ levels in exuded sap reflect the O_2_ available to planktonic bacteria *in planta*, such bulk analyses can mask the spatial heterogeneity within plant hosts. Additionally, we recently used X-ray microtomography to show that phyl. I strain GMI1000 induces wilt symptoms when its dense biofilms clog half of the total xylem vessels (4). The SEM images shown here suggest that phyl. II/IV strains may form biofilms that have more available oxygen.

It is surprising that although all RSSC strains experience low oxygen conditions *in planta*, phyl. II/IV strains benefit little from the presence of the alternate TEA NO_3_^-^. We previously hypothesized that RSSC strains may have an O_2_ and NO_3_^-^-independent mechanism to obtain energy in VDM, such as amino acid fermentation or Stickland reactions (3). However, Stickland metabolism is rare outside of *Clostridia* (45), and our data are consistent with the conclusion that RSSC lack fermentative metabolism and are obligate O_2_ and NO_3_^-^respirors. These bacteria may have such a high affinity for oxygen that they can scavenge the extremely small amount of oxygen in the medium during growth in an anaerobic chamber.

Differences in denitrification metabolism correlated with phenotypic differences in aerotaxis behavior and biofilm formation. When allowed to migrate to their preferred oxygen concentration in soft agar, phyl. I/III strains moved to lower O_2_ levels in a NO_3_^-^dependent manner. Without NO_3_^-^, these strains preferred the O_2-_rich agar surface. When NO_3_^-^was present, they migrated to a deeper, less oxygen-rich band. In contrast, the phyl. II/IV strains were indifferent to the presence of NO_3_^-^. Among the RSSC, aerotaxis has only been studied in phyl. II strain K60 (33).

Using a population genetics test, several denitrification-related genes were identified as core RSSC genes under selection (46). The Tajima’s D were above 2 for the accessory metabolic proteins NarI and NarJ and the NO_3_^-^-responsive regulators NarL and NarX, suggesting that there could be functionally distinct alleles of these genes within the RSSC population. Except for a robust study of variation in quorum sensing synthases/regulators within the RSSC (47), there has been relatively little investigation of variation of regulators within the RSSC.

In addition to differences in denitrifying metabolism, our genomic enrichment analyses revealed that phyl. I/III and phyl. II/IV vary in their predicted capacity for degrading aromatic compounds. Aromatic compounds are a major class of plant defense chemicals as well as possible carbon sources. Several aromatics have been detected and quantified in tomato xylem sap during infection with phyl. I strain GMI1000: salicylic acid (~20-200 nM), benzoic acid (~10 nM), and coumaric acid (~100 nM) (48). Our analyses found that phyl. II/IV genomes were differentially enriched in a benzoic acid/catechol degradation pathway (29). The ability to degrade other aromatic compounds significantly increases *in planta* fitness of phyl. I strain GMI1000 (31, 32). It remains to be determined whether benzoate/catechol degradation similarly contributes to *in planta* fitness of phyl. II/IV strains.

Across the domain Bacteria, genes encoding energy metabolism are among those most commonly found in recently horizontally acquired regions (49). The evolution of the RSSC has clearly been shaped by HGT events, likely boosted by the group’s natural competency (50, 51). Strains in all RSSC phylotypes take up and transfer DNA (51), although phyl. I is the most recombinogenic of the phylotypes (51–54). The saprophytic life stages of phyl. I/III strains may have provided opportunities to horizontally acquire the *nos* cluster from other soil residents. Moreover, survival in the soil may exert selective pressure for complete denitrification. Like the RSSC, *Bradyrhizobium* isolates vary in the presence/absence of the *nosZ* gene. An elegant microbial ecology study in Japan found that *nosZ*-minus *Bradyrhizobium* strains dominated in soil types that have high levels of volcanic ash (55). In contrast, *Bradyrhizobium* strains with the full denitrification pathway have higher tolerance to flooding (56, 57). Little is known about whether RSSC lineages vary in their ability to survive in different soil types, but some lineages persist better in soil than others (58, 59).

## CONCLUSIONS

Our genomic, physiological, and virulence studies collectively suggest that RSSC strains in phyl. I/III and II/IV use different metabolic strategies to reach high cell densities *in planta*. Phyl. I/III strains benefit from denitrifying respiration during tomato infection, including N_2_O reduction by NosZ. However, phyl. II/IV strains are fully virulent, growing to similar densities in host stems and causing identical wilt symptoms even though they lack NosZ and cannot complete denitrifying respiration. Broader genomic and behavioral analyses *in planta* and *in vitro* suggest that the two groups respond differently to oxygen, a key environmental variable. The completely denitrifying strains in phyl. I/III grow better under hypoxic conditions *in vitro* and are more likely to aggregate in biofilms on host plant xylem vessel walls. Phyl. II/IV strains have respiratory strategies that allow them to exploit environmental and host-associated niches containing higher oxygen levels. Ongoing studies will test the intriguing hypothesis that these divergent energy strategies reflect inter-species niche partitioning.

## MATERIALS AND METHODS

### Strains, mutagenesis, and culture conditions

Origins and accession numbers of strains used in this study are in Table S1. One representative strain for each RSSC phylotype was selected, based on ability to cause disease on the common host tomato, and on existing experimental data and closed genome sequences for strains GMI1000, CMR15, K60, and PSI07. The Δ*nosZ* GMI1000 mutant was generated using SOE-PCR to replace the *nosZ* ORF with a gentamicin resistance gene cassette as described (3). This mutation was moved into strain CMR15 with natural transformation (50). All strains were maintained in −80°C glycerol stocks and cultured on solid CPG plates prior to growth in broth.

Bacterial growth was measured in modified VDM medium (0.5 g KH_2_PO_4_, 0.5 g K_2_HPO_4_, 0.234 g MgSO_4_, 2.5 g casamino acids, 50 mM sodium succinate, +/-30 mM potassium nitrate) at 28°C under controlled O_2_ conditions (3, 60). Bacteria were routinely cultured in rich CPG broth. Unless otherwise noted, media were inoculated to a starting O.D._600_ of 0.001 (~1×10^6^ CFU/mL) and endpoint data were collected 72 h post inoculation. Anaerobic growth was in BD GasPak systems in 1.7 mL tubes; O.D._600_ was measured spectrophotometrically at the endpoint. Aerobic assays were incubated in a 28°C shaker at 225 rpm in 96-well plates sealed with Breathe-Easy membrane (Sigma-Aldrich). All other O_2_ conditions were generated in a gas-controlled chamber (Ruskinn Invivo_2_400, Sanford, ME) in 96-well plates sealed with breathable tape. For 96-well plate assays, a Synergy HT microtiter plate reader (Biotek Instruments, Winooski VT) was used to measure endpoint A_600_.

To assess bacterial growth under a range of O_2_ conditions, we stab-inoculated semi-solid VDM (0.2% noble agar) with ~1×10^7^ CFU of each strain. Tubes were incubated at 28°C without shaking and visually assessed after a week. NO_2_^-^ was quantified (reported as +/-) using Griess reactions (Molecular Probes, Inc. Eugene, OR) in lysed cell supernatant of anaerobic cultures following 3 h incubation at high cell densities (O.D._600_ = 1.0; 1×10^9^ CFU/mL) as described (3). N_2_ production was visually assessed as presence of gas bubbles over 96 h of anaerobic incubation in 1.7 mL tubes following inoculation at O.D._600_ = 0.001. All assays were replicated 3 times per treatment per strain.

### N_2_O quantification

Overnight aerobic cultures of representative strains from each phylotype were diluted to an O.D.600 of 0.100 (~1×10^8^ CFU/ml). 100 μl of culture was added to 100 ml of VDM media containing 10 mM NO_3_^-^in a 300 ml flask. For these assays NO_3_^-^concentration was reduced to 10 mM to avoid saturating the instrument. Flasks were sealed with a double-holed rubber-stopper with two glass tubes inserted for flushing the headspace and sampling gas. Flasks were flushed with >3 volumes of N_2_ gas to create anaerobic conditions and glass tubes were stopped with small rubber septa. Media was not deoxygenated, because RSSC cultures rapidly deoxygenate media biologically (3). Cultures were incubated statically at 28 °C for 24 h, at which point 100 μl of culture was removed and dilution plated to enumerate CFU/ml. Nitrous oxide gas generated by denitrification was measured using a needle and syringe to draw out a 300 mL gas sample from the top of the flask, while N_2_ gas replaced withdrawn gas just above the culture level. Gas samples were placed in GC vials, injected into an Agilent 7890A GC System (Santa Clara, CA, USA) and analyzed as described (61).

### Growth in host tomato plants and virulence assays

To measure pathogen growth *in planta*, 21 day-old tomato plants (wilt-susceptible cv. Bonny Best) were inoculated through a cut leaf petiole with 2000 CFU of RSSC as described (3). Two days post-inoculation, 0.1 g stem tissue was collected from the midstem directly above the inoculated petiole, ground, and dilution plated to quantify bacterial population size. These assays contained 10 plants per strain.

To assess relative virulence, unwounded 21-day-old tomato plants were soil-soak inoculated with 1×10^8^ CFU of RSSC per g potting mix as described (3). Symptoms were assessed daily using a disease index based on percent of leaves wilted (0=healthy, 1=1-25%, 2=26-50%, 3=51-75%, 4=76-100%) (62). Each assay contained 16 plants per treatment, and the assays were replicated three times.

### Phylogenetic analysis

We selected 51 genomes that represent the genomic diversity of the RSSC (63). Using KBase (64), we built a phylogenetic tree by using the “Insert Set of Genomes into Species Tree” app, which creates an MSA from 49 well-conserved bacterial proteins and builds a tree with FastTree2 version 2.1.10 (65).

To expand the analysis beyond the RSSC, all complete genomes of Betaproteobacteria publicly available on NCBI at the time of this analysis (2012) were compared using MLST based on 31 loci as described (66). The ORFcore pipeline (67) and MUSCLE 3.7 (68) were used to extract, correct, align, and concatenate amino acid sequences. Fasttree v2.1.3 was used for tree construction (65). A gene was considered present if it shared >40% (or in the case of NosF >30%) amino acid identity over at least 70% of the length of the query sequence (*Pseudomonas stutzeri* CAA37) as determined with blastp in the BLAST+ package. Blastp was also used for amino acid comparisons across all available NosZ sequences on NCBI at the time of analysis (2015).

### Protein sequence comparisons

Homologs of denitrification pathway genes were detected in genomes of strains GMI1000, CMR15, K60 and PSI07 using the MicroScope web interface, BLAST, and OMA (69). The corresponding protein sequences were aligned using MUSCLE (68) and percentage identity to strain GMI1000 was computed for each locus using the seqinr R package (70).

### FNR binding site predictions

The Virtual Footprint Regulon Analysis program from PRODORIC predicted intergenic FNR binding sites across RSSC□genomes□using a weighted matrix generated from the *E. coli* FNR binding sequence (71). Sequence logos were generated with□Weblogo (72), and the trimmed FNR binding site sequences from all four phylotypes. □The PRODORIC Virtual Footprint Promoter Analysis program predicted the presence of FNR binding sites upstream of genes related to denitrification for three representative strains. Because its genome was then in draft form, the representative Phyl. II strain K60 was excluded from this analysis.

### KEGG enrichment analysis

All RSSC genome sequences available in 2015 were annotated using HMMer models for KEGG and Pfam as described (73). Amino acid identity (25% with 50% coverage) was used to cluster sequences *de novo* with proteinorthov5. Gene families over-represented in Phyl. I/III or Phyl. II/IV strains were identified using Fisher’s exact test in the Python package scipy (74). The significance level was set at *P*>0.05.

### Biofilm assays

Biofilm formation of each phylotype representative strain was assessed *in vitro* using the PVC plate-crystal violet stain assay (33). Values were reported as O.D._590_/O.D._600_ to normalize for minor differences in growth rates among strains.

### Scanning electron microscopy

Three-week-old Bonny Best tomato plants were inoculated through a cut leaf petiole with 1×10^2^ CFU with RSSC GMI1000, K60, CMR15, or PSI07. At first sign of symptoms (DI=1), two thin slices were collected from each mid-stem following surface sterilization and processed for SEM as described (35). Samples were visualized with a Zeiss LEO 1530 high-resolution scanning electron microscope (Materials Sciences Center, University of Wisconsin-Madison). For each strain, 6-8 plants were assessed.

### Bacterial streaming assay

Nineteen-day-old tomato plants were inoculated through a cut petiole with 1×10^3^ CFU of strain GMI1000, K60, CMR15, or PSI07 as described above. At the first sign of symptoms (DI=1), 1 cm of stem tissue from 0.5 cm below the inoculation site was excised, cut in half, and floated in 1 ml sterile water in a 24-well microplate. Samples were incubated for 90 min at room temperature with 85 rpm shaking before the stem section was removed and the escaped bacteria were quantified as OD_600 nm_ using a spectrophotometer.

### Xylem sap oxygen measurement

Seventeen-day-old tomato plants were soil soak inoculated as described above with 1×10^8^ CFU/ml of either GMI1000, CMR15, K60, PSI07, or water. When symptoms first appeared, disease severity was rated, plants were de-topped and xylem sap rapidly pooled on the cut stem via natural root pressure. An oxygen microsensor probe (Unisense, Aarhus, Denmark) was immediately inserted into the pooled xylem sap and read until the signal was steady for 60 s. For an O_2_ saturated water control, air was bubbled through distilled water for 5 min. The anoxic control solution was 0.1 M sodium ascorbate, 0.1 M sodium hydroxide in water. The O_2_ content of stagnant deionized water was measured at each sampling point to ensure probe consistency.

## ACKNOWLEDGEMENTS

We gratefully acknowledge Mick Blue, Robert Anex, and Ian Rowland for HPLC and NMR analysis. Additionally, we thank Matthew Pereyra for SEM technical support. We thank Valley Stewart, Becky Parales, and many Allen lab members for helpful comments and suggestions.

## Supplementary Materials

**Table S1.** *Ralstonia* spp. strains used in this study.

| Strain | Phyl | Host | Origin | Accession Number |
| --- | --- | --- | --- | --- |
| **GMI1000** | **I** | **Tomato** | **Guyana** | **Genbank: NC_003295, NC_003296** |
| Y45 | I | Tobacco | China | GenBank: AFWL000000000 |
| F504, FQY_4 | I | Tobacco | China | GenBank: NC_020799.1, 021745.1 |
| YC45 | I | Ginger | China | GenBank: CP011997.1, CP011998.1 |
| Rs09-161 | I | Eggplant | India | GenBank: CM002757.1, CM002758.1 |
| Rs10-244 | I | Pepper | India | GenBank: CM002755.1, CM002756.1 |
| R292 | I | Mulberry | China | NA |
| PSS81 | I | Tomato | Taiwan | NA |
| **K60** | **II** | **Tomato** | **United States** | **GenBank: NCTK000000000** |
| B50 | II | Banana | Brazil | EMBL: PRJEB7421 |
| IBSBF1900 | II | Banana | Brazil | EMBL: PRJEB8309 |
| CFBP2957 | II | Tomato | French West Indies | EMBL: FP885897, FP885907 |
| UW181 | II | Plantain | Venezuela | EMBL: PRJEB8309 |
| Grenada9-1 | II | Banana | Grenada | EMBL: PRJEB7428 |
| Po82 | II | Potato | Mexico | GenBank: CP002819, CP002820 |
| UW179 | II | Banana | Colombia | EMBL: PRJEB7426 |
| UW163 | II | Plantain | Peru | EMBL: PRJEB7430 |
| IBSBF1503 | II | Cucumber | Brazil | EMBL: PRJEB7433 |
| CFBP6783 | II | Heliconia | French West Indies | EMBL: PRJEB7432 |
| CFBP7014 | II | Anthurium | Trinidad | EMBL: PRJEB8309 |
| CFBP1416 | II | Plantain | Costa Rica | EMBL: PRJEB7434 |
| CIP417 | II | Banana | Philippines | EMBL: PRJEB7437 |
| MolK2 | II | Banana | Phillipines | GenBank: CAHW01000040 |
| CFBP3858 | II | Potato | Netherlands | EMBL: PRJEB8309 |
| UW349 | II | Potato | Brazil | GenBank: JQOI000000000.1 |
| UW365 | II | Potato | China | GenBank: JQSI000000000.1 |
| IPO1609 | II | Potato | Netherlands | GenBank: CU914168, CUP914166 |
| UW551 | II | Geranium | Kenya | GenBank: AAKL000000000 |
| UY031 | II | NA | Brazil | GenBank: CP012687.1, CP012688.1 |
| UW491 | II | Potato | Colombia | GenBank: JQSH000000000.1 |
| RS2 | II | Potato | NA | EMBL: PRJEB8309 |
| **CMR15** | **III** | **Tomato** | **Cameroon** | **EMBL: FP885895, FP885896** |
| J25 | III | Tomato | Kenya | NA |
| CFBP3059 | III | Eggplant | Burkinia Faso | NA |
| **PSI07** | **IV** | **Tomato** | **Indonesia** | **EMBL: FP885906, FP885891** |
| R229 | IV | Banana | Indonesia | EMBL: FP854059 to FR854085 |
| KACC10722 | IV | NA | South Korea | GenBank: CP014702.1, CP014703.1 |
| R24 | IV | Clove | Indonesia | EMBL: FR854086 to FR854092 |
| CFBP6727 | IV | Potato | Indonesia | NA |
| MAFF301558 | IV | Potato | Japan | NA |

NA = not available

**Figure S1.**
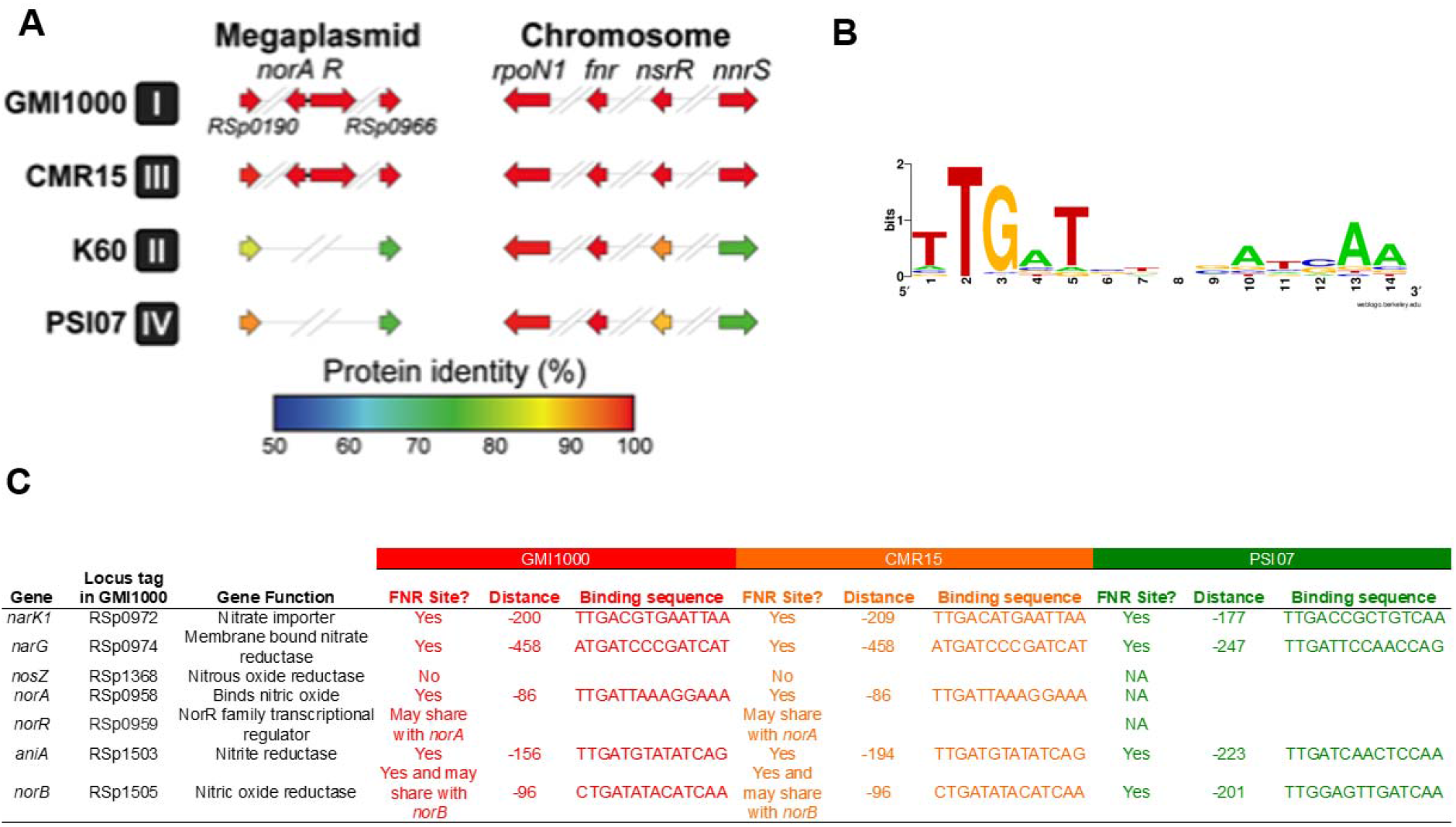
Denitrification-related regulators and FNR predicted binding sites in genomes of phylotype representative strains in the RSSC. **A)** The presence or absence and % amino acid identity of all denitrification-associated regulators were determined, using the strain GMI1000 genome for comparison. The *norAR* gene pair is predicted to be nitric oxide-responsive, while *rpoN1* controls gene expression in response to nitrogen starvation and *nsrR* responds to nitrite/nitric oxide. NnrS in other systems responds to nitric oxide. The majority of the regulators, particularly FNR-like RSp0190, RSp0966 and *fnr* (RSc1283), are predicted to be oxygen-responsive. **B)** Predicted binding site consensus (logo) for the RSSC FNR-like regulon. **C)** Predicted FNR-like regulator binding sites located upstream of start site for inorganic nitrogen metabolism genes in strains GMI1000, CMR15, and PSI07.

**Figure S2.**
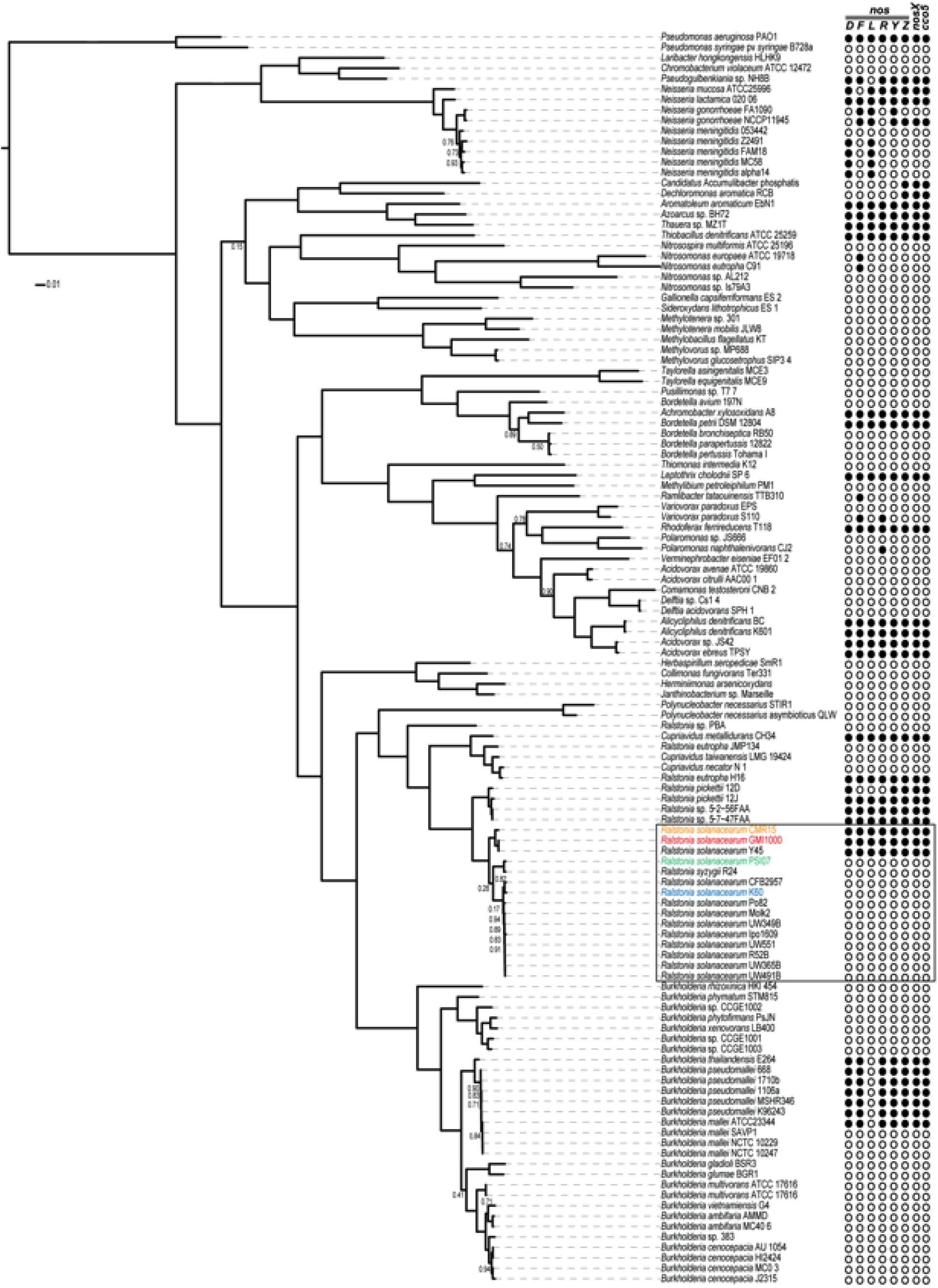
The *nos* cluster, containing the genes required for the last step of denitrification, has been repeatedly laterally transferred across the Betaproteobacteria. All complete Betaproteobacteria genomes from NCBI available in 2012 were analyzed. Nos protein sequences were identified using a strict BlastP threshold for annotation (>40% identity over 70% of the query sequence length for all but NosF, which had a cut-off of 30% because it is more diverse). *P. stutzeri* CAA37 Nos sequences were used as the query sequences using blastp in the BLAST+ package. Circles represent the 6 standard components of the nitrous oxide reductase gene cluster (listed in alphabetical order from *nosD* to *nosZ*) and 2 additional genes (*nosX* and *cco5*) that often co-segregate. White circles indicate absence of a gene and black circles indicate its presence. All strains in the RSSC are framed in the black rectangle. Strains chosen to represent each of the four phylotypes are in color: phylotype I representative, GMI1000, is in red; phylotype II K60 in blue; phylotype III CMR15 in orange; phylotype IV PSI07 in green.

**Figure S3.**
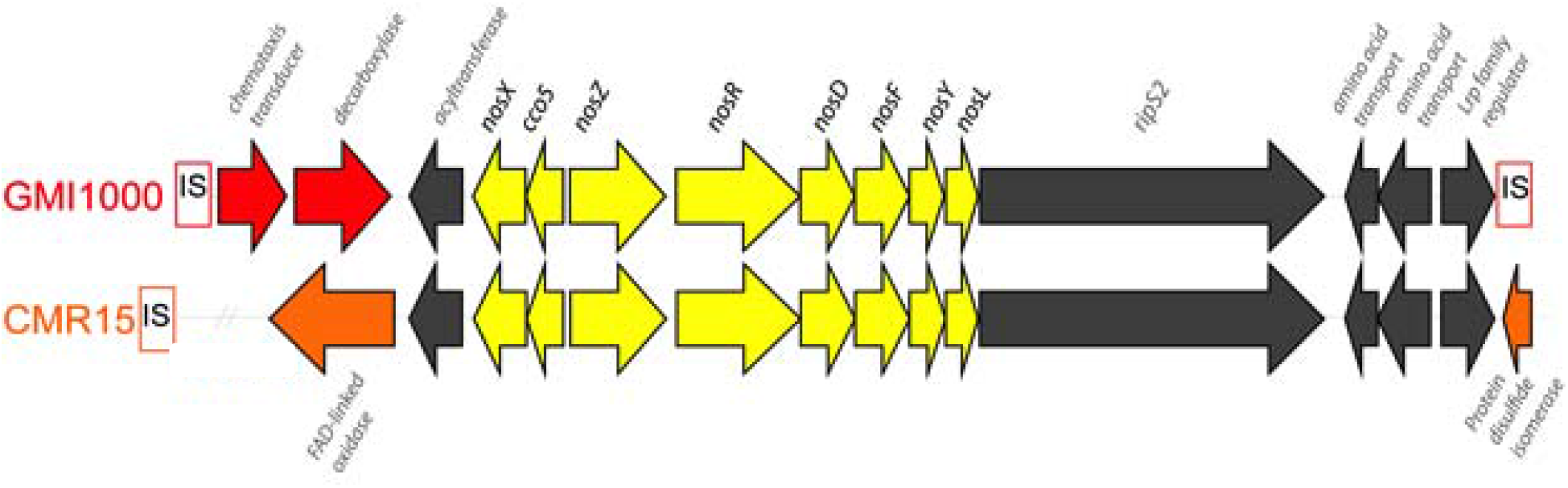
*R. solanacearum* strains in phylotypes I and III share a highly conserved nitrous oxide reductase gene cluster (in yellow). All genes near the *nosZ* ORF that lie between two predicted insertion sequences in phyl. I strain GMI1000 genome (locus tags RSp1362-RSp1378) are aligned to the region in phyl. III strain CMR15 (CMR15_mp30001-CMR15_mp30026). Genes found only in GMI1000 are in red while genes in orange are specific to CMR15. Predicted functions based on NCBI and Phyre2 protein structure modeling are listed in gray italics near each ORF lacking an annotation.

**Figure S4.**
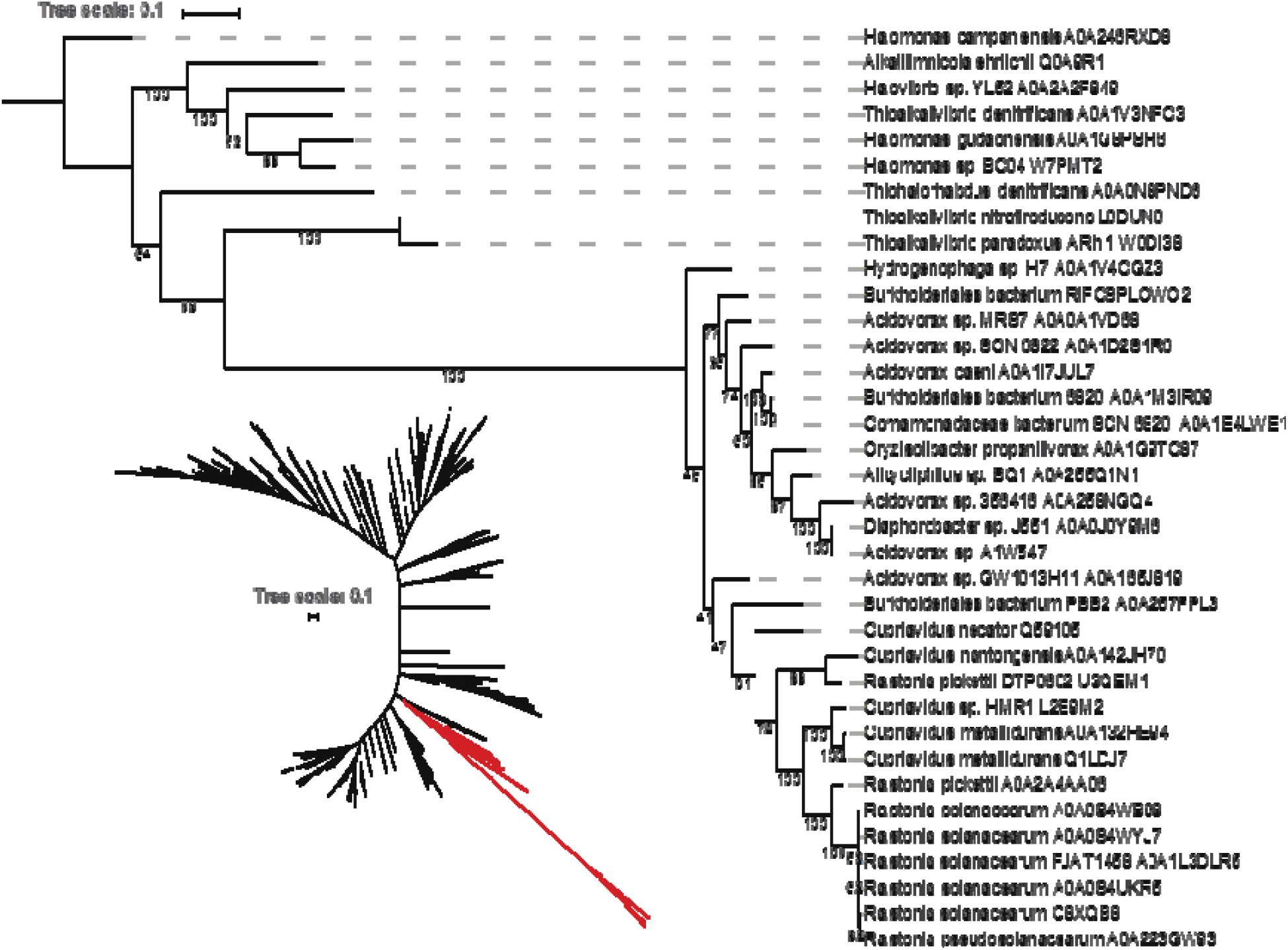
NosZ protein phylogeny. Inset: Full phylogeny of all NosZ protein sequences from uniprot. The clade containing *Ralstonia* is highlighted in red. The clade containing Uniprot proteins from the RSSC is shown in detail; they form a monophyletic clade of highly related sequences.

**Figure S5.**
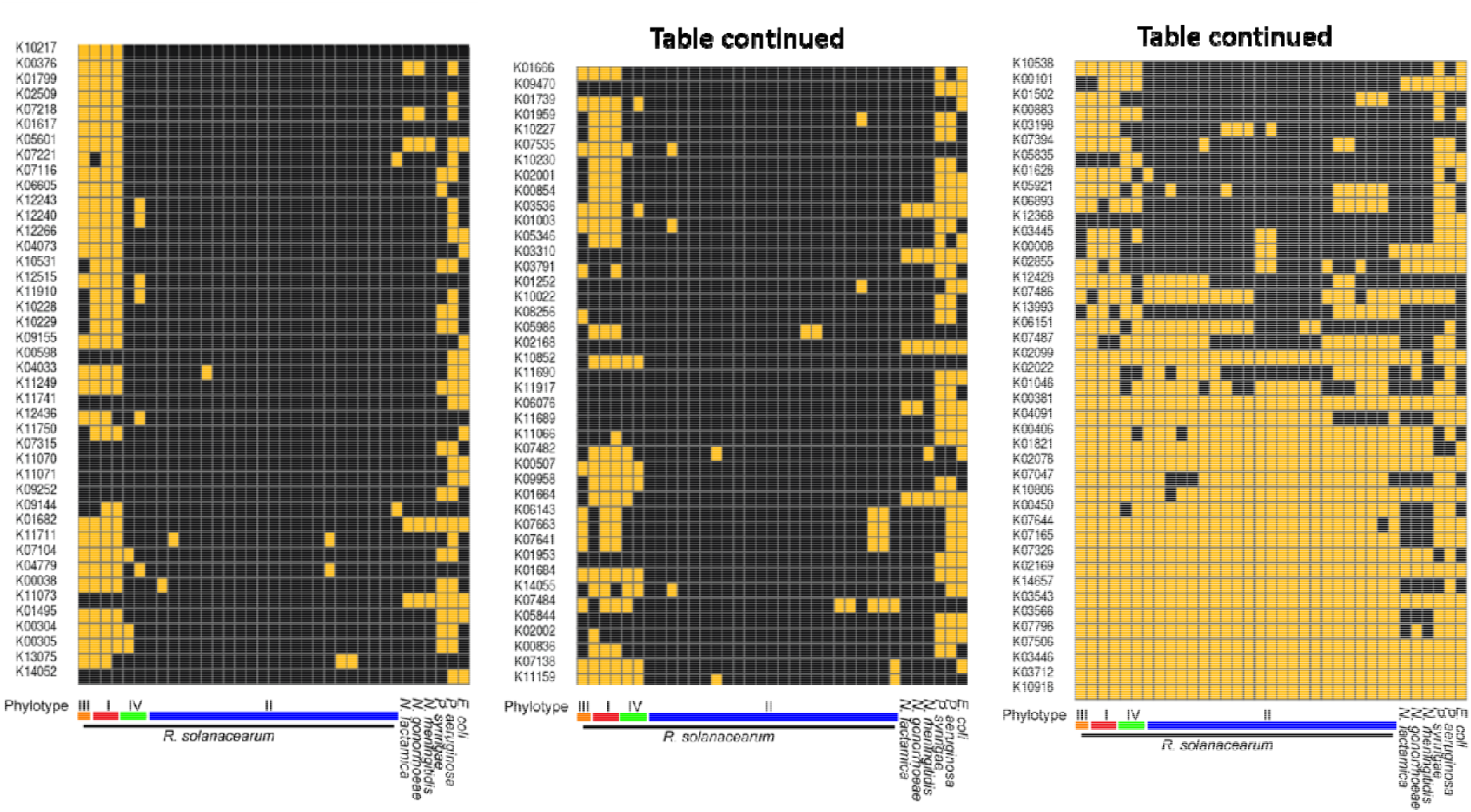
Over-represented KEGG groups in RSSC phylotype I and III strains. HMMer models for KEGG were used to identify the over-represented KEGG families in *R. solanacearum*. Complete denitrifiers I and III shown in red and orange respectively, while incomplete denitrifiers II and IV are shown in blue and green respectively. Presence/absence of overrepresented KEGG families depicted above as yellow or black respectively. Distantly related bacteria with and without complete denitrification were used for comparison (right side of figure).

**Figure S6.**
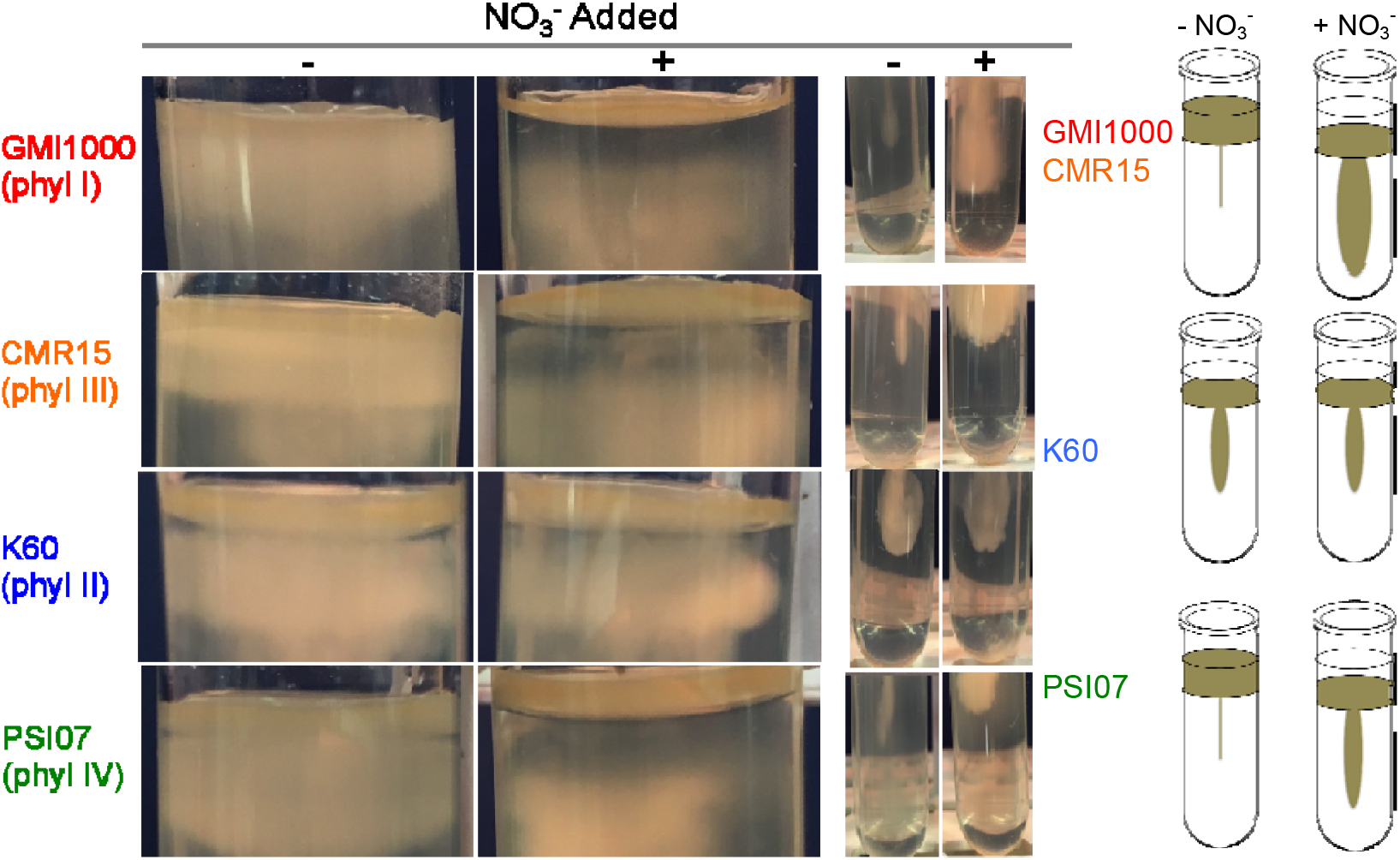
Oxygen preference assessments of *R. solanacearum* phylotype representatives. CPG overnight cultures of each strain were pelleted and re-suspended in water to an O.D._600_ of 1.0. 10uL of these cell suspensions were stab inoculated and slowly released by a pipette into a plastic tube filled with 20 mL 0.2% semisolid VDM agar with or without nitrate added. Following one week of 28°C incubation without shaking, tubes were imaged. To the right, diagrams depict the general trends.

**Figure S7.**
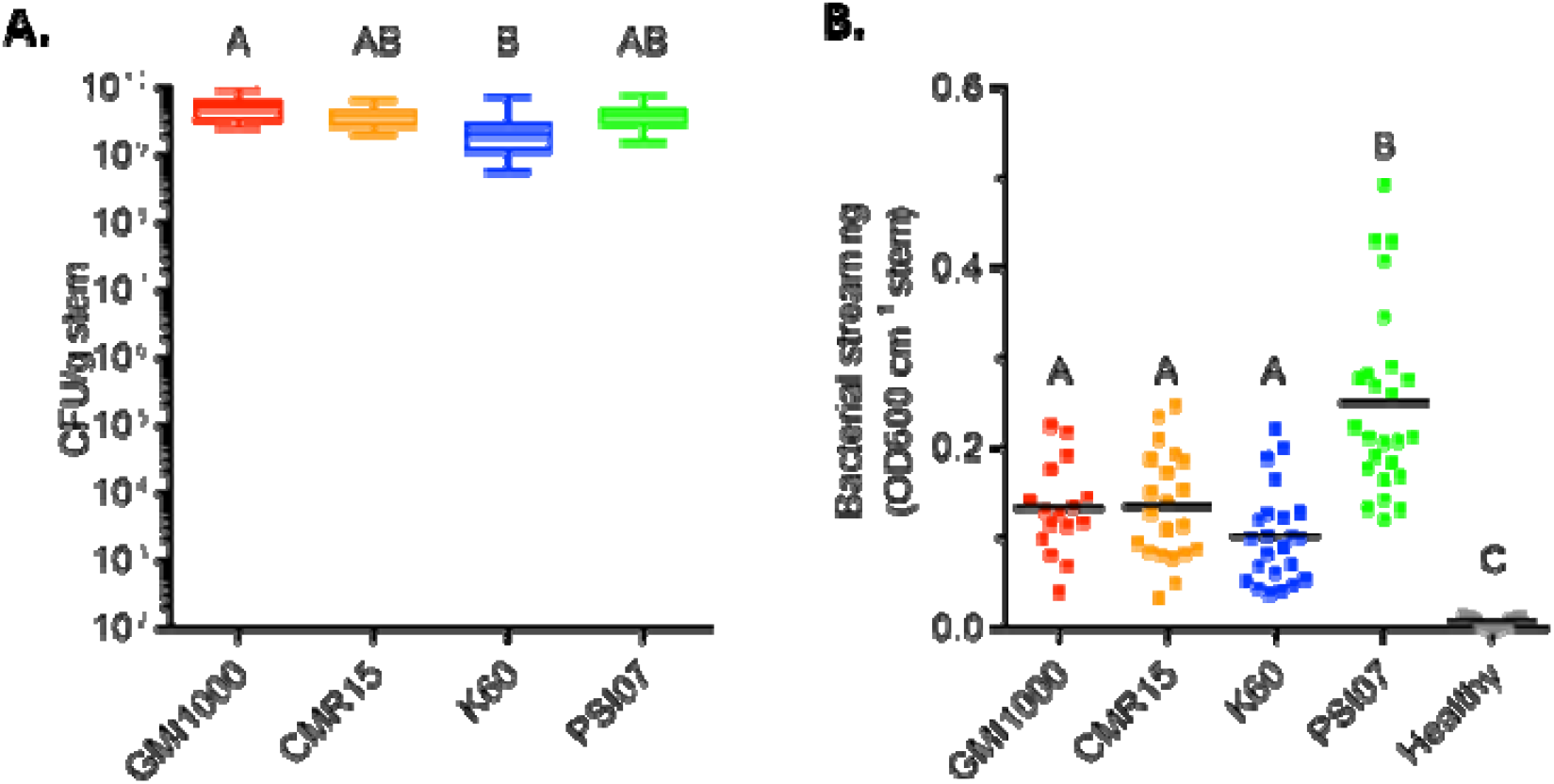
Complete and incomplete RSSC denitrifiers adhere to xylem differently. Representative tomato-colonizing strains (GMI1000, K60, CMR15, PSI07) were inoculated into tomato stem via cut petiole inoculation (1×10^3^ CFU). At the first sign of wilting symptoms, **(A)** total bacterial populations in the stem were enumerated by dilution plating or **(B)** bacterial attachment to stem xylem was assessed by measuring planktonic cells that floated or swam out of cut stem tissue. One cm of stem was incubated in water with 85 rpm shaking. After 90 min, bacterial density in the water was measured by OD_600 nm_. Letters indicate *P*<0.05 by ANOVA with Tukey’s multiple comparison test; (A) N=14-15 plants and (B) N=16-25 plants per condition.

## REFERENCES

1. Fischbach MA, Sonnenburg JL. 2011. Eating for two: how metabolism establishes interspecies interactions in the gut. Cell Host Microbe 10:336–347.

2. Zumft WG. 1997. Cell biology and molecular basis of denitrification. Microbiol Mol Biol Rev 61:533–616.

3. Dalsing BL, Truchon AN, Gonzalez-Orta ET, Milling AS, Allen C. 2015. *Ralstonia solanacearum* uses inorganic nitrogen metabolism for virulence, ATP production, and detoxification in the oxygen-limited host xylem environment. MBio 6:e02471.

4. Ingel B, Caldwell D, Duong F, Parkinson DY, McCulloh KA, Iyer-Pascuzzi AS, McElrone AJ, Lowe-Power TM. 2022. Revisiting the source of wilt symptoms: X-ray microcomputed tomography provides direct evidence that *Ralstonia* biomass clogs xylem vessels. Phytofrontiers https://doi.org/10.1094/PHYTOFR-06-21-0041-R.

5. Prior P, Ailloud F, Dalsing BL, Remenant B, Sanchez B, Allen C. 2016. Genomic and proteomic evidence supporting the division of the plant pathogen *Ralstonia solanacearum* into three species. BMC Genomics 17:90.

6. Remenant B, Coupat-Goutaland B, Guidot A, Cellier G, Wicker E, Allen C, Fegan M, Pruvost O, Elbaz M, Calteau A, Salvignol G, Mornico D, Mangenot S, Barbe V, Médigue C, Prior P. 2010. Genomes of three tomato pathogens within the *Ralstonia solanacearum* species complex reveal significant evolutionary divergence. BMC Genomics 11:379.

7. Safni I, Cleenwerck I, De Vos P, Fegan M, Sly L, Kappler U. 2014. Polyphasic taxonomic revision of the *Ralstonia solanacearum* species complex: proposal to emend the descriptions of *R. solanacearum* and *R. syzygii* and reclassify current *R. syzygii* strains. Int J Syst Evol Microbiol 2014;64:3087–103.

8. Genin S, Denny TP. 2012. Pathogenomics of the Ralstonia solanacearum species complex. Annu Rev Phytopathol 50:67–89.

9. Lowe-Power TM, Khokhani D, Allen C. 2018. How *Ralstonia solanacearum* exploits and thrives in the flowing plant xylem environment. Trends Microbiol 26:929–942.

10. Peyraud R, Cottret L, Marmiesse L, Gouzy J, Genin S. 2016. A Resource Allocation Trade-Off between Virulence and Proliferation Drives Metabolic Versatility in the Plant Pathogen Ralstonia solanacearum. PLoS Pathog 12:e1005939.

11. Hamilton CD, Steidl OR, MacIntyre AM, Hendrich CG, Allen C. 2021. Ralstonia solanacearum Depends on Catabolism of Myo-Inositol, Sucrose, and Trehalose for Virulence in an Infection Stage–Dependent Manner. Mol Plant Microbe Interact 34:669–679.

12. Gerlin L, Escourrou A, Cassan C, Maviane Macia F, Peeters N, Genin S, Baroukh C. 2021. Unravelling physiological signatures of tomato bacterial wilt and xylem metabolites exploited by Ralstonia solanacearum. Environ Microbiol 23:5962–5978.

13. Baroukh C, Zemouri M, Genin S. 2022. Trophic preferences of the pathogen Ralstonia solanacearum and consequences on its growth in xylem sap. Microbiologyopen 11:e1240.

14. Plener L, Boistard P, González A, Boucher C, Genin S. 2012. Metabolic adaptation of Ralstonia solanacearum during plant infection: a methionine biosynthesis case study. PLoS One 7:e36877.

15. Táncsics, Farkas, Szoboszlay, Szabó. 2013. One-year monitoring of meta-cleavage dioxygenase gene expression and microbial community dynamics reveals the relevance of subfamily I. 2. C extradiol … Syst Acarol Acarol Spec Publ.

16. Nestler H, Kiesel B, Kaschabek SR, Mau M, Schlömann M, Balcke GU. 2007. Biodegradation of chlorobenzene under hypoxic and mixed hypoxic-denitrifying conditions. Biodegradation 18:755–767.

17. Remenant B, de Cambiaire J-C, Cellier G, Jacobs JM, Mangenot S, Barbe V, Lajus A, Vallenet D, Medigue C, Fegan M, Allen C, Prior P. 2011. *Ralstonia syzygii,* the Blood Disease Bacterium and some Asian *R. solanacearum* strains form a single genomic species despite divergent lifestyles. PLoS One 6:e24356.

18. Rodionov DA, Dubchak IL, Arkin AP, Alm EJ, Gelfand MS. 2005. Dissimilatory metabolism of nitrogen oxides in bacteria: comparative reconstruction of transcriptional networks. PLoS Comput Biol 1:e55.

19. Ray SK, Kumar R, Peeters N, Boucher C, Genin S. 2015. rpoN1, but not rpoN2, is required for twitching motility, natural competence, growth on nitrate, and virulence of Ralstonia solanacearum. Front Microbiol 6:229.

20. Truchon AN, Hendrich CG, Bigott AF, Dalsing BL, Allen C. 2022. NorA, HmpX, and NorB Cooperate to Reduce NO Toxicity during Denitrification and Plant Pathogenesis in Ralstonia solanacearum. Microbiol Spectr 10:e0026422.

21. Barth KR, Isabella VM, Clark VL. 2009. Biochemical and genomic analysis of the denitrification pathway within the genus Neisseria. Microbiology 155:4093–4103.

22. Holloway P, McCormick W, Watson RJ, Chan YK. 1996. Identification and analysis of the dissimilatory nitrous oxide reduction genes, nosRZDFY, of Rhizobium meliloti. J Bacteriol 178:1505–1514.

23. Cuypers H, Viebrock-Sambale A, Zumft WG. 1992. NosR, a membrane-bound regulatory component necessary for expression of nitrous oxide reductase in denitrifying Pseudomonas stutzeri. J Bacteriol 174:5332–5339.

24. Dell’acqua S, Moura I, Moura JJG, Pauleta SR. 2011. The electron transfer complex between nitrous oxide reductase and its electron donors. J Biol Inorg Chem 16:1241–1254.

25. Saunders NF, Hornberg JJ, Reijnders WN, Westerhoff HV, de Vries S, van Spanning RJ. 2000. The NosX and NirX proteins of Paracoccus denitrificans are functional homologues: their role in maturation of nitrous oxide reductase. J Bacteriol 182:5211–5217.

26. Wunsch P, Körner H, Neese F, van Spanning RJM, Kroneck PMH, Zumft WG. 2005. NosX function connects to nitrous oxide (N2O) reduction by affecting the CuZ center of NosZ and its activity in vivo. FEBS Lett 579:4605–4609.

27. Wunsch P, Herb M, Wieland H, Schiek UM, Zumft WG. 2003. Requirements for Cu(A) and Cu-S center assembly of nitrous oxide reductase deduced from complete periplasmic enzyme maturation in the nondenitrifier Pseudomonas putida. J Bacteriol 185:887–896.

28. Jacobs JM, Babujee L, Meng F, Milling A, Allen C. 2012. The *in planta* transcriptome of *Ralstonia solanacearum:* conserved physiological and virulence strategies during bacterial wilt of tomato. MBio 3.

29. Harwood CS, Parales RE. 1996. The beta-ketoadipate pathway and the biology of self-identity. Annu Rev Microbiol 50:553–590.

30. Powlowski J, Shingler V. 1994. Genetics and biochemistry of phenol degradation byPseudomonas sp. CF600. Biodegradation 5:219–236.

31. Lowe TM, Ailloud F, Allen C. 2015. Hydroxycinnamic acid degradation, a broadly conserved trait, protects *Ralstonia solanacearum* from chemical plant defenses and contributes to root colonization and virulence. Mol Plant Microbe Interact 28:286–297.

32. Lowe-Power TM, Jacobs JM, Ailloud F, Fochs B, Prior P, Allen C. 2016. Degradation of the plant defense signal salicylic acid protects *Ralstonia solanacearum* from toxicity and enhances virulence on tobacco. MBio 7.

33. Yao J, Allen C. 2007. The plant pathogen *Ralstonia solanacearum* needs aerotaxis for normal biofilm formation and interactions with its tomato host. J Bacteriol 189:6415–6424.

34. Stewart PS, Franklin MJ. 2008. Physiological heterogeneity in biofilms. Nat Rev Microbiol 6:199–210.

35. Minh Tran T, MacIntyre A, Khokhani D, Hawes M, Allen C. 2016. Extracellular DNases of *Ralstonia solanacearum* modulate biofilms and facilitate bacterial wilt virulence. Environ Microbiol 18:4103–4117.

36. Khokhani D, Lowe-Power TM, Tran TM, Allen C. 2017. A single regulator mediates strategic switching between attachment/spread and growth/virulence in the plant pathogen *Ralstonia solanacearum*. mBio 8.

37. Georgoulis S, Shalvarjian KE, Helmann TC, Hamilton CD, Carlson HK, Deutschbauer AM, Lowe-Power TM. 2021. Genome-Wide Identification of Tomato Xylem Sap Fitness Factors for Three Plant-Pathogenic *Ralstonia* Species. mSystems 6:e01229–21.

38. Dalsing BL, Allen C. 2014. Nitrate assimilation contributes to *Ralstonia solanacearum* root attachment, stem colonization, and virulence. J Bacteriol 196:949–960.

39. Hendrich CG, Truchon AN, Dalsing BL, Allen C. 2021. Nitric Oxide Regulates the *Ralstonia solanacearum* Type 3 Secretion System. bioRxiv.

40. Flores-Cruz Z, Allen C. 2009. *Ralstonia solanacearum* encounters an oxidative environment during tomato infection. Mol Plant Microbe Interact 22:773–782.

41. Flores-Cruz Z, Allen C. 2011. Necessity of OxyR for the hydrogen peroxide stress response and full virulence in *Ralstonia solanacearum*. Appl Environ Microbiol 77:6426–6432.

42. Colburn-Clifford JM, Scherf JM, Allen C. 2010. *Ralstonia solanacearum* Dps contributes to oxidative stress tolerance and to colonization of and virulence on tomato plants. Appl Environ Microbiol 76:7392–7399.

43. Colburn-Clifford J, Allen C. 2010. A cbb3-type cytochrome C oxidase contributes to Ralstonia solanacearum R3bv2 growth in microaerobic environments and to bacterial wilt disease development in tomato. Mol Plant Microbe Interact 23:1042–1052.

44. Siletsky SA, Borisov VB. 2021. Proton Pumping and Non-Pumping Terminal Respiratory Oxidases: Active Sites Intermediates of These Molecular Machines and Their Derivatives. Int J Mol Sci 22.

45. Pavao A, Graham M, Arrieta-Ortiz ML, Immanuel SRC, Baliga NS, Bry L. 2022. Reconsidering the in vivo functions of Clostridial Stickland amino acid fermentations. Anaerobe 76:102600.

46. Eckshtain-Levi N, Weisberg AJ, Vinatzer BA. 2018. The population genetic test Tajima’s D identifies genes encoding pathogen-associated molecular patterns and other virulence-related genes in *Ralstonia solanacearum*. Mol Plant Pathol 19:2187–2192.

47. Kai K, Ohnishi H, Shimatani M, Ishikawa S, Mori Y, Kiba A, Ohnishi K, Tabuchi M, Hikichi Y. 2015. Methyl 3-hydroxymyristate, a diffusible signal mediating *phc* quorum sensing in *Ralstonia solanacearum*. Chembiochem 16:2309–2318.

48. MacIntyre AM, Meline V, Gorman Z, Augustine SP, Dye CJ, Hamilton CD, Iyer-Pascuzzi AS, Kolomiets MV, McCulloh KA, Allen C. 2022. Trehalose increases tomato drought tolerance, induces defenses, and increases resistance to bacterial wilt disease. PLoS One 17:e0266254.

49. Nakamura Y, Itoh T, Matsuda H, Gojobori T. 2004. Biased biological functions of horizontally transferred genes in prokaryotic genomes. Nat Genet 36:760–766.

50. Bertolla F, Van Gijsegem F, Nesme X, Simonet P. 1997. Conditions for natural transformation of Ralstonia solanacearum. Appl Environ Microbiol 63:4965–4968.

51. Guidot A, Coupat B, Fall S, Prior P, Bertolla F. 2009. Horizontal gene transfer between Ralstonia solanacearum strains detected by comparative genomic hybridization on microarrays. ISME J 3:549–562.

52. Wicker E, Lefeuvre P, de Cambiaire J-C, Lemaire C, Poussier S, Prior P. 2012. Contrasting recombination patterns and demographic histories of the plant pathogen Ralstonia solanacearum inferred from MLSA. ISME J 6:961–974.

53. Prokchorchik M, Pandey A, Moon H, Kim W, Jeon H, Jung G, Jayaraman J, Poole S, Segonzac C, Sohn KH, McCann HC. 2020. Host adaptation and microbial competition drive *Ralstonia solanacearum* phylotype I evolution in the Republic of Korea. Microb Genom 6.

54. Sharma P, Johnson MA, Mazloom R, Allen C, Heath LS, Lowe-Power TM, Vinatzer BA. 2022. Meta-analysis of the *Ralstonia solanacearum* species complex (RSSC) based on comparative evolutionary genomics and reverse ecology. Microb Genom 8.

55. Shiina Y, Itakura M, Choi H, Saeki Y, Hayatsu M, Minamisawa K. 2014. Relationship between soil type and N□O reductase genotype (nosZ) of indigenous soybean bradyrhizobia: nosZ-minus populations are dominant in Andosols. Microbes Environ 29:420–426.

56. Saeki Y, Nakamura M, Mason MLT, Yano T, Shiro S, Sameshima-Saito R, Itakura M, Minamisawa K, Yamamoto A. 2017. Effect of Flooding and the *nosZ* Gene in Bradyrhizobia on Bradyrhizobial Community Structure in the Soil. Microbes Environ 32:154–163.

57. Salas A, Tortosa G, Hidalgo-García A, Delgado A, Bedmar EJ, Richardson DJ, Gates AJ, Delgado MJ. 2019. The Hemoglobin Bjgb From Bradyrhizobium diazoefficiens Controls NO Homeostasis in Soybean Nodules to Protect Symbiotic Nitrogen Fixation. Front Microbiol 10:2915.

58. Ray JD, Subandiyah S, Prakoso AB, Rincón-Flórez VA, Carvalhais LC, Drenth A. 2022. Transmission of Blood Disease in Banana. Plant Dis 106:2155–2164.

59. Sequeira L. 1993. Bacterial wilt: past, present, and future, p. 12–12. In Aciar Proceedings. AUSTRALIAN CENTRE FOR INTERNATIONAL AGRICULTURAL RESEARCH.

60. Hayward AC, El-Nashaar HM, Nvdegger U, Lindo LD. 1990. Variation in nitrate metabolism in biovars of Pseudomonas solanacearum. J Appl Bacteriol 69:269–280.

61. Collier SM, Ruark MD, Oates LG, Jokela WE, Dell CJ. 2014. Measurement of greenhouse gas flux from agricultural soils using static chambers. J Vis Exp e52110.

62. Khokhani D, Tran TM, Lowe-Power TM, Allen C. 2018. Plant assays for quantifying *Ralstonia solanacearum* virulence. Bio-protocol.

63. Lowe-Power T, Avalos J, Charco Munoz M, Chipman K. 2020. A meta-analysis of the known global distribution and host range of the *Ralstonia* species complex. bioRxiv.

64. Arkin AP, Cottingham RW, Henry CS, Harris NL, Stevens RL, Maslov S, Dehal P, Ware D, Perez F, Canon S, Sneddon MW, Henderson ML, Riehl WJ, Murphy-Olson D, Chan SY, Kamimura RT, Kumari S, Drake MM, Brettin TS, Glass EM, Chivian D, Gunter D, Weston DJ, Allen BH, Baumohl J, Best AA, Bowen B, Brenner SE, Bun CC, Chandonia J-M, Chia J-M, Colasanti R, Conrad N, Davis JJ, Davison BH, DeJongh M, Devoid S, Dietrich E, Dubchak I, Edirisinghe JN, Fang G, Faria JP, Frybarger PM, Gerlach W, Gerstein M, Greiner A, Gurtowski J, Haun HL, He F, Jain R, Joachimiak MP, Keegan KP, Kondo S, Kumar V, Land ML, Meyer F, Mills M, Novichkov PS, Oh T, Olsen GJ, Olson R, Parrello B, Pasternak S, Pearson E, Poon SS, Price GA, Ramakrishnan S, Ranjan P, Ronald PC, Schatz MC, Seaver SMD, Shukla M, Sutormin RA, Syed MH, Thomason J, Tintle NL, Wang D, Xia F, Yoo H, Yoo S, Yu D. 2018. KBase: The United States Department of Energy Systems Biology Knowledgebase. Nat Biotechnol 36:566–569.

65. Price MN, Dehal PS, Arkin AP. 2010. FastTree 2--approximately maximum-likelihood trees for large alignments. PLoS One 5:e9490.

66. Ciccarelli FD, Doerks T, von Mering C, Creevey CJ, Snel B, Bork P. 2006. Toward automatic reconstruction of a highly resolved tree of life. Science 311:1283–1287.

67. Klassen JL, Currie CR. 2013. ORFcor: identifying and accommodating ORF prediction inconsistencies for phylogenetic analysis. PLoS One 8:e58387.

68. Edgar RC. 2004. MUSCLE: multiple sequence alignment with high accuracy and high throughput. Nucleic Acids Res 32:1792–1797.

69. Altenhoff AM, Škunca N, Glover N, Train C-M, Sueki A, Piližota I, Gori K, Tomiczek B, Müller S, Redestig H, Gonnet GH, Dessimoz C. 2015. The OMA orthology database in 2015: function predictions, better plant support, synteny view and other improvements. Nucleic Acids Res 43:D240–9.

70. Charif D, Lobry JR. 2007. SeqinR 1.0-2: A Contributed Package to the R Project for Statistical Computing Devoted to Biological Sequences Retrieval and Analysis, p. 207–232. In Bastolla, U, Porto, M, Roman, HE, Vendruscolo, M (eds.), Structural Approaches to Sequence Evolution: Molecules, Networks, Populations. Springer Berlin Heidelberg, Berlin, Heidelberg.

71. Münch R, Hiller K, Barg H, Heldt D, Linz S, Wingender E, Jahn D. 2003. PRODORIC: prokaryotic database of gene regulation. Nucleic Acids Res 31:266–269.

72. Crooks GE, Hon G, Chandonia J-M, Brenner SE. 2004. WebLogo: a sequence logo generator. Genome Res 14:1188–1190.

73. Murfin KE, Lee M-M, Klassen JL, McDonald BR, Larget B, Forst S, Stock SP, Currie CR, Goodrich-Blair H. 2015. Xenorhabdus bovienii Strain Diversity Impacts Coevolution and Symbiotic Maintenance with Steinernema spp. Nematode Hosts. MBio 6:e00076.

74. Virtanen P, Gommers R, Oliphant TE, Haberland M, Reddy T, Cournapeau D, Burovski E, Peterson P, Weckesser W, Bright J, van der Walt SJ, Brett M, Wilson J, Millman KJ, Mayorov N, Nelson ARJ, Jones E, Kern R, Larson E, Carey CJ, Polat İ, Feng Y, Moore EW, VanderPlas J, Laxalde D, Perktold J, Cimrman R, Henriksen I, Quintero EA, Harris CR, Archibald AM, Ribeiro AH, Pedregosa F, van Mulbregt P, SciPy 1.0 Contributors. 2020. SciPy 1.0: fundamental algorithms for scientific computing in Python. Nat Methods 17:261–272.

